# Disease-relevant single cell photonic signatures identify S100β stem cells and their myogenic progeny in vascular lesions

**DOI:** 10.1101/2020.05.13.093518

**Authors:** Claire Molony, Damien King, Mariana Di Luca, Michael Kitching, Abidemi Olayinka, Roya Hakimjavadi, Lourdes A.N. Julius, Emma Fitzpatrick, Yusof Gusti, Denise Burtenshaw, Killian Healy, Emma K. Finlay, David Kernan, Andreu Llobera, Weimin Liu, David Morrow, Eileen M. Redmond, Jens Ducrée, Paul A. Cahill

**Affiliations:** Vascular Biology and Therapeutics Laboratory, School of Biotechnology Faculty of Science and Health, Dublin City University, Dublin 9, Ireland; Fraunhofer Project Centre for Embedded BioAnalytical Systems, Faculty of Science and Health, Dublin City University, Dublin 9, Ireland; School of Biotechnology Faculty of Science and Health, Dublin City University, Dublin 9, Ireland; Centre Nacional de Microelectronica, Campus UAB, Barcelona, Spain; Department of Surgery, University of Rochester Medical Center, Rochester, NY USA

**Keywords:** photonics, autofluorescence imaging, S100β vascular stem cells, carotid artery ligation, lineage tracing, arteriosclerosis, smooth muscle differentiation, multivariate analysis, supervised machine learning

## Abstract

A hallmark of subclinical atherosclerosis is the accumulation of vascular smooth muscle cell (SMC)-like cells leading to intimal thickening and lesion formation. While medial SMCs contribute to vascular lesions, the involvement of resident vascular stem cells (vSCs) remains unclear. We evaluated single cell photonics as a discriminator of cell phenotype *in vitro* before the presence of vSC within vascular lesions was assessed *ex vivo* using supervised machine learning and further validated using lineage tracing analysis. Using a novel lab-on-a-Disk (Load) platform, label-free single cell photonic emissions from normal and injured vessels *ex vivo* were interrogated and compared to freshly isolated aortic SMCs, cultured Movas SMCs, macrophages, B-cells, S100β^+^ mVSc, bone marrow derived mesenchymal stem cells (MSC) and their respective myogenic progeny across five broadband light wavelengths (λ465 – λ670 ± 20 nm). We found that profiles were of sufficient coverage, specificity, and quality to clearly distinguish medial SMCs from different vascular beds (carotid vs aorta), discriminate normal carotid medial SMCs from lesional SMC-like cells *ex vivo* following flow restriction, and identify SMC differentiation of a series of multipotent stem cells following treatment with transforming growth factor beta 1 (TGF-β1), the Notch ligand Jagged1, and Sonic Hedgehog using multivariate analysis, in part, due to photonic emissions from enhanced collagen III and elastin expression. Supervised machine learning supported genetic lineage tracing analysis of S100β^+^ vSCs and identified the presence of S100β^+^ vSC-derived myogenic progeny within vascular lesions. We conclude disease-relevant photonic signatures may have predictive value for vascular disease.

## Introduction

Cardiovascular disease (CVD), the leading cause of death and disability world-wide, is characterized by pathological structural changes to the blood vessel wall [1]. A hallmark is the accumulation of smooth muscle cell (SMC)-like cells leading to intimal medial thickening (IMT) and the obstruction to blood flow that may culminate in a heart attack or stroke [2], Observations in humans vessels confirm that early ‘transitional’ lesions enriched with SMC-like cells are routinely present in atherosclerotic-prone regions of arteries during pathologic intimal thickening, lipid retention, and the appearance of a developed plaque [3]. Endothelial cell (EC) dysfunction due to disturbed blood flow is classically associated with the development of atheroprone lesions [4] and is routinely modelled in mouse carotid arteries following flow restriction caused by ligation [5]. These lesions can further develop into advanced atherosclerotic plaques in *ApoE* knockout mice on Western diets [6].

Label-free technologies for bio-analytical applications have attracted recent attention. Several different classes of label-free sensor detectors are being developed including plasmonic, electrical, mechanical, and photonic (light) sensors [7], Light as a diagnostic and prognostic tool has several potential advantages including high sensitivity, non-destructive measurement, non-invasive analysis and low limits of detection [7], The innate optical response comprises scattering, absorbance, and auto-fluorescence signals that are closely associated with metabolism [8] and structural proteins [9] under normal, altered or pathological conditions, while fluorescence has been proposed as a non-invasive indicator of embryonic stem cell differentiation state [10]. In combination with microfluidics, photonics enables real time measurement of single cells in very small sample volumes [11]. In this context, we have developed label-free photonic technologies with highly efficient cell-to-light coupling that have been successfully deployed to measure the real-time response of individual cells in a population [12], To accompany these photonic platforms, deep learning, a subset of machine learning based primarily on artificial neural network geometries, has rapidly grown as a predictive method to obtain real-time bio-photonic decision-making systems and analyze bio-photonic data, in particular, spectroscopic data including spectral data pre-processing and spectral classification [13].

Several key modulators of vascular SMC phenotype and fate have been reported. Transforming growth factor (TGF-β1) is a cytokine regulating myogenic differentiation that is released by both inflammatory cells and vascular cells [endothelial (EC) and SMC] [14, 15] within the vessel wall and has been implicated in the etiology of atherosclerosis [16], vessel restenosis [17], and the development of the neointima [18]. Similarly, Notch and Hedgehog signalling components are secreted as pro-atherogenic stimuli to promote lesion formation [19–22],

Herein, we utilised a novel LoaD platform to measure label-free single cell photonic emissions across five broadband light wavelengths (λ465 – λ670 ± 20 nm) from normal and injured vessels *ex vivo* and compared them to freshly isolated aortic SMCs, cultured Movas SMCs, macrophages, B-cells, S100β^+^ resident mVSc, and bone marrow derived mesenchymal stem cells (MSC) and their respective myogenic progeny. We then combined single cell photonics with supervised machine learning using multilayer perceptron (MLP) neural network analysis and linear discriminant analysis (LDA) of these trained datasets, in conjunction with lineage tracing of S100β^+^ mVSc *in vivo*, to examine the cellular heterogeneity of vascular lesions following iatrogenic flow restriction.

## Results

### Single cell photonics of cells from normal and injured vessels *ex vivo*

Flow restriction following carotid artery ligation resulted in a significant increase in adventitial and intimal volumes after 14 days when compared to sham control [Figure 1A, B] concomitant with a significant reduction in the number of Myh11^+^ positive cells and expression of Myh11 in protein lysates [Figure 1C, D], In contrast, medial and intimal SMC-like cells maintained expression of smooth muscle cell α-actin (SMA) [Figure 1C]. Medial cells were isolated and single cell photonic profiles of individual cells *ex vivo* were measured following capture on V-cups using a centrifugal Lab-on-a-Disc (LoaD) platform, as previously described [12] [Figure 1E], and visualised by phase contrast microscopy on each V-cup [Figure 1F], The Log10 autofluorescence emissions of 356 cells from 6 animals/group were recorded across five broadband light wavelengths (λ465, λ530, λ565, λ630 and λ670 with a bandwidth of 20 nm) [Figure 1G] and corrected as Log2 fold increase over background [Figure 1H], The photonic intensity of single cells from the ligated vessels *ex vivo* was significantly enhanced at the first 3 wavelengths (λ465, λ530 and λ565) when compared to single cells from sham-operated control vessels [Figure 1G, H], Principal component analysis (PCA) revealed that the photonic profile at λ565 had a significant influence on the PC1 components [Figure 1I] with significant divergence from other variables suggesting that this variable was poorly correlated with other wavelengths [Figure 1J], Linear discriminant analysis (LDA) revealed that the photonic profile of cells from sham and ligated vessels could be easily separated from each other across the five wavelengths [Figure 1K], Confusion matrices revealed that 88% of the sham cells were classified as similar to each other with only a small proportion of cells classified as similar to cells from ligated vessels [Figure 1L], In contrast, 21% of ligated cells were classified as similar to sham cells with an accuracy of 86.2% on a cross-validated leave-one-out basis [Figure 1L],

**Figure 1.**
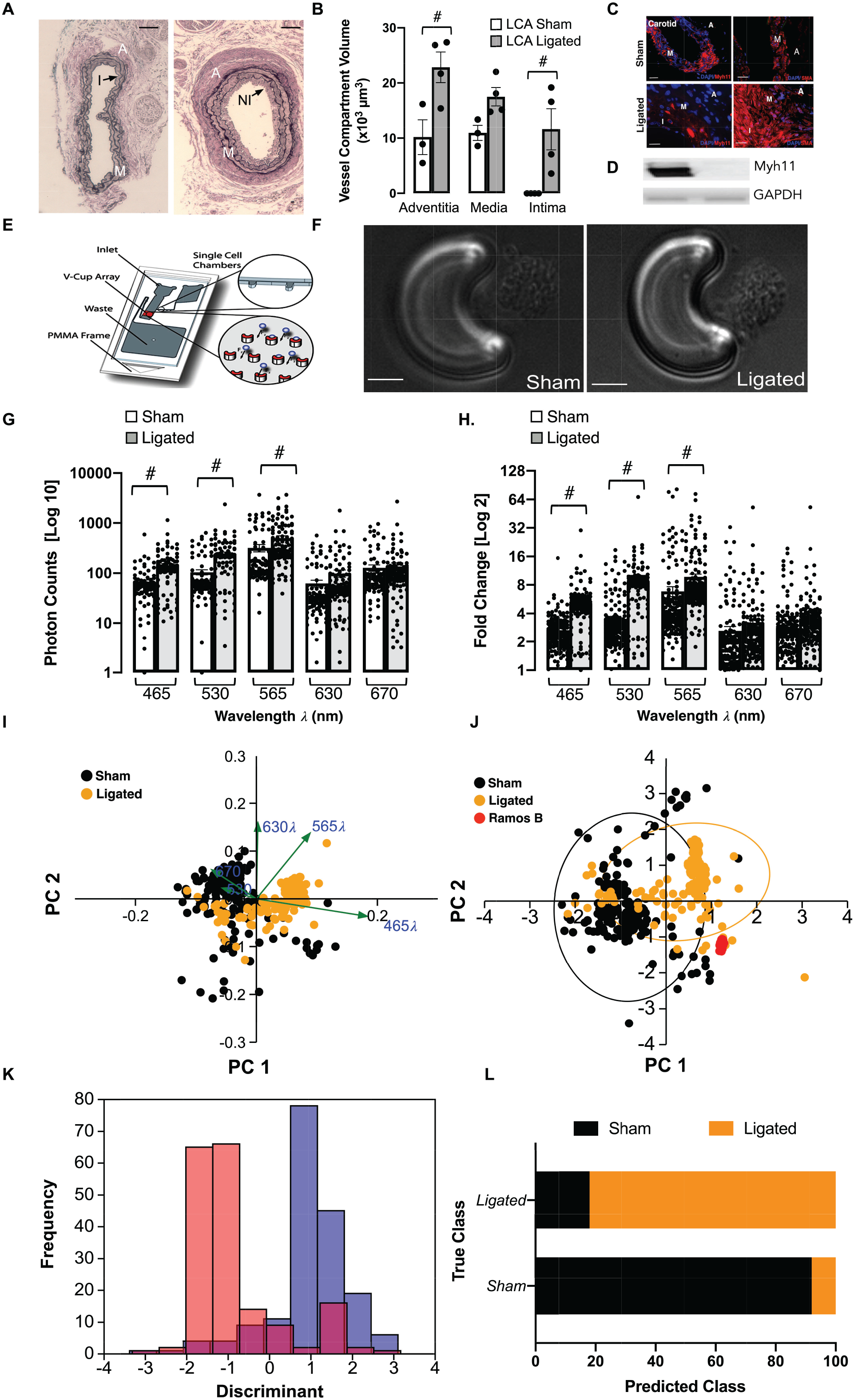
Single cell photonics from normal and injured vessels *ex vivo* following flow restriction. **A**. Representative haematoxylin and eosin (H&E) staining of LCA from sham and ligated vessels 14 days post-ligation (A: Adventitia, M: Media, I: Intima and NI:Neointima). **B**. Morphometric analysis of adventitial, medial and intimal volumes within the LCA in sham and ligated arteries; (#p≤0.05, n=3). C. Representative immunohistochemical analysis of Myh11 (red) and α-actin expression (red) and DAPI nuclei staining (blue) in adventitial, medial and intimal layers of the LCA from sham and ligated carotid arteries. **D**. Representative immunoblot of Myh11 expression in the LCA from sham and ligated carotid artery (n=3). E. Schematic of capture of individual cells using the LoaD platform. F. Representative image of formalin-fixed cells from sham and ligated vessels captured on a V-cup using the LoaD platform. **G**. Single cell auto-fluorescence photon emissions across five broadband light wavelengths (λ465, λ530, λ565, λ630 and λ670 nm with a bandwidth of 20 nm) from sham and ligated LCA *ex vivo*. Data are the mean ± SEM of Log10 photons from 178 cells/group from 6 pooled vessels, #p≤0.001 vs sham. H. Log2 fold increase over background of single cell photon emissions from sham and ligated LCA. Data are the mean ± SEM of 178 cells/group from 6 pooled vessels, #p≤0.001 vs sham. I. PCA biplots of single cell photon analysis from sham (black) and ligated (orange) vessels. Each *λ* variable is representative of the eigenvectors (loadings). J. PCA loading plots of single cell photon analysis from sham (black) and ligated (orange) cells *ex vivo* compared to Ramos B cells *in vitro* (red). Data are from 411 cells across the five wavelengths. K. LDA plots of single cell photons from sham (black) and ligated (orange) vessels. L. Confusion matrix of true class representing the given group assignment and predicted class representing the estimated group assignment following a leave-one-out cross-validation procedure.

These data indicate that single-cell autofluorescence emissions across five wavelengths *ex vivo* are of sufficient coverage, specificity and overall quality to unambiguously identify the majority of carotid artery cells from sham vessels as differentiated Myh11^+^, SMA^+^ contractile SMCs. In contrast, the majority of cells from injured vessels exhibit a distinct spectral profile with a minority of cells similar to differentiated SMCs from sham animals *ex vivo*.

### Single cell photonics of B-cells and macrophages *in vitro*

To confirm that photonics can discriminate disparate cell phenotypes, single cell photonic measurements were recorded in human B lymphocytes (Ramos B) and murine macrophages (J774A.1) across the same five wavelengths and visualised by phase contrast microscopy on each V-cup [Suppl Figure 1A]. Multivariate analysis demonstrated that single cells clustered tightly and away from each other [Suppl Figure 1B] while both cell types could be easily separated from each other [Suppl Figure 1C]. When compared to cells from sham and ligated vessels [Suppl Figure 1D-G], Ramos B and J774A.1 cells clustered away from these cells and were easily separated from each other [Suppl Figure 1H-J]. These data indicate that single-cell photonics clearly identifies lymphocytes and macrophages as distinct cell populations compared to isolated cells of sham and ligated vessels

### Single cell photonics of aortic SMCs ex vivo and cultured SMCs

To test whether differentiated SMCs differ in their photonic profile from de-differentiated SMCs in culture, freshly isolated mouse aortic SMCs *ex vivo* and sub-cultured Movas SMCs *in vitro* were visualised by phase contrast microscopy on each V-cup [Figure 2A] before single cell photonic emissions were recorded. Both cell types had similar photonic intensities across most wavelengths [Figure 2B] but had lower photonic intensities when compared to Ramos B cells [Figure 2C] and were clearly separated by LDA on a cross-validated leave-one-out basis with 87.8% accuracy [Figure 2D], Multivariate analysis demonstrated that a fraction of Movas SMC clustered towards aortic SMCs with a proportion of Movas SMC classified similar to aortic SMCs [Figure 2E],

**Figure 2.**
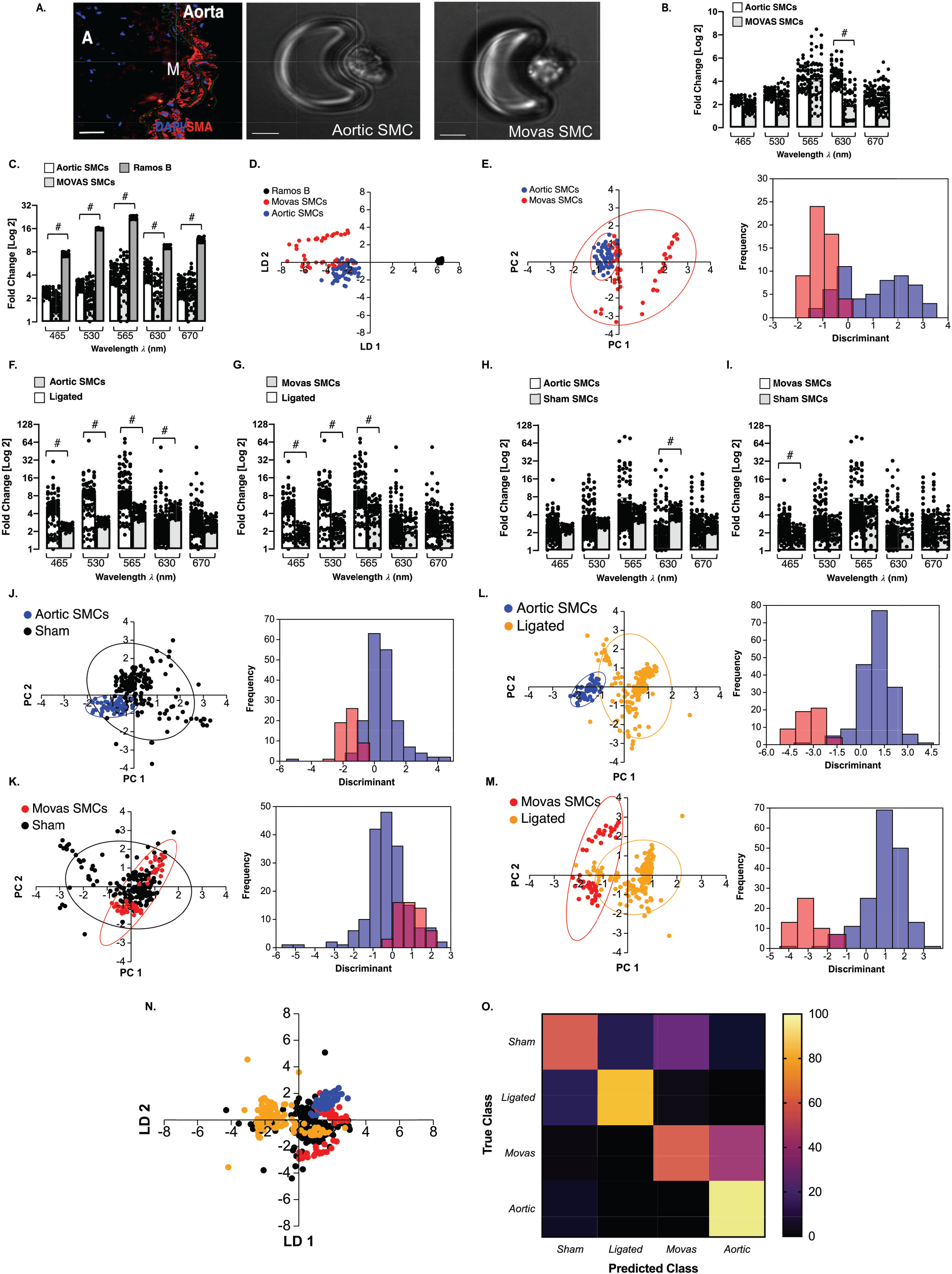
Single cell photonics of aortic SMCs *ex vivo* and cultures aortic Movas SMCs *in vitro*. **A**. Representative immunohistochemical analysis of ⍰-actin (SMA) expression in mouse aorta and visualisation of aortic SMCs and Movas SMCs on each V-cup in the LoaD platform. B. Single cell photon emissions across five wavelengths from freshly isolated aortic SMCs and cultured aortic Movas SMCs *in vitro*. Data are the Log2 fold increase and represent the mean ± SEM of 55 cells/group, #p≤0.001. C. Single cell photon emissions of aortic and Movas SMCs compared to Ramos B cells. Data are the Log2 fold increase and represent the mean ± SEM of 55 cells/group, #p≤0.001. **D**. LDA plots of aortic SMC (blue), Movas SMC (red) and Ramos B (black) cells *in vitro*. Data are from 165 cells across the five wavelengths E. PCA loading and LDA plots of aortic SMC and Movas SMC. **F-I**. Single cell photon emissions from (F, H) aortic SMC and (G, I) Movas SMC cells compared to sham (F, G) and ligated cells (H, I) *ex vivo*. Data are the Log2 fold increase and represent the mean ± SEM of 55-178 cells/group, #p≤0.001 vs Sham (F,G) Ligated (H,I). **J-M**. PCA loading plots and LDA of aortic SMCs (J, L) and Movas SMCs (K, M) compared to carotid artery SMCs (sham) (J) and ligated cells, respectively. **N**. LDA plots of sham (black), ligated (orange) cells *ex vivo* compared to aortic SMCs (blue) and Movas SMC (Red) cells *in vitro* **G**. Confusion matrix of true class and predicted class following a leave-one-out cross-validation procedure by the LDA classifier. Data are from 466 cells across five wavelengths.

To determine if differentiated SMCs differ between vascular beds (aortic vs carotid), the photonic profile of freshly isolated aortic SMCs and Movas SMC were compared to normal and ligated carotid artery cells *ex vivo*. Aortic SMCs and Movas SMC had similar photonic intensities to carotid artery SMCs across most wavelengths [Figure 2F, G] but exhibited lower photonic intensities compared to cells of ligated vessels [Figure 2H, I], Multivariate analysis revealed that aortic SMCs and Movas SMC were distinct to carotid artery SMCs [Figure 2J, K] and ligated cells [Figure 2L, M], Moreover, while a proportion of cells isolated from sham vessels were classified as similar to Movas SMCs (26%) and aortic SMCs (6%), respectively [Figure 2N], the majority of cells from ligated vessels were distinct with an accuracy of 72% on a cross-validated leave-one-out basis [Figure 2O].

### Single cell photonics of bone marrow-derived mesenchymal stem cells and their myogenic progeny

Single cell photonics of undifferentiated bone marrow-derived mesenchymal stem cells (MSCs) were compared to their myogenic SMC progeny following stimulation with media supplemented with TGF-β1. Myogenic differentiation was confirmed by epigenetic, biochemical and immunohistochemical analysis of cells before and after treatment with TGF-β1 [Figure 3A-D], Bone marrow-derived MSCs treated with TGF-β1 (2ng/mL) for 7 days increased their expression of SMC differentiation markers calponin 1 (Cnn1) and myosin heavy chain 11 (Myh11) [Figure 3A, B], concomitant with a significant increase in the enrichment of the SMC epigenetic histone mark di-methylation of lysine 3 on Histone 4 (H3K4me2) [23] and a decrease in the repressive mark tri-methylation of lysine 27 on Histone 3 (H3K27me3)[24] at the Myh11 locus [Figure 3C], and an increase in Cnn1 and Myh11 expression in protein lysates [Figure 3D],

**Figure 3.**
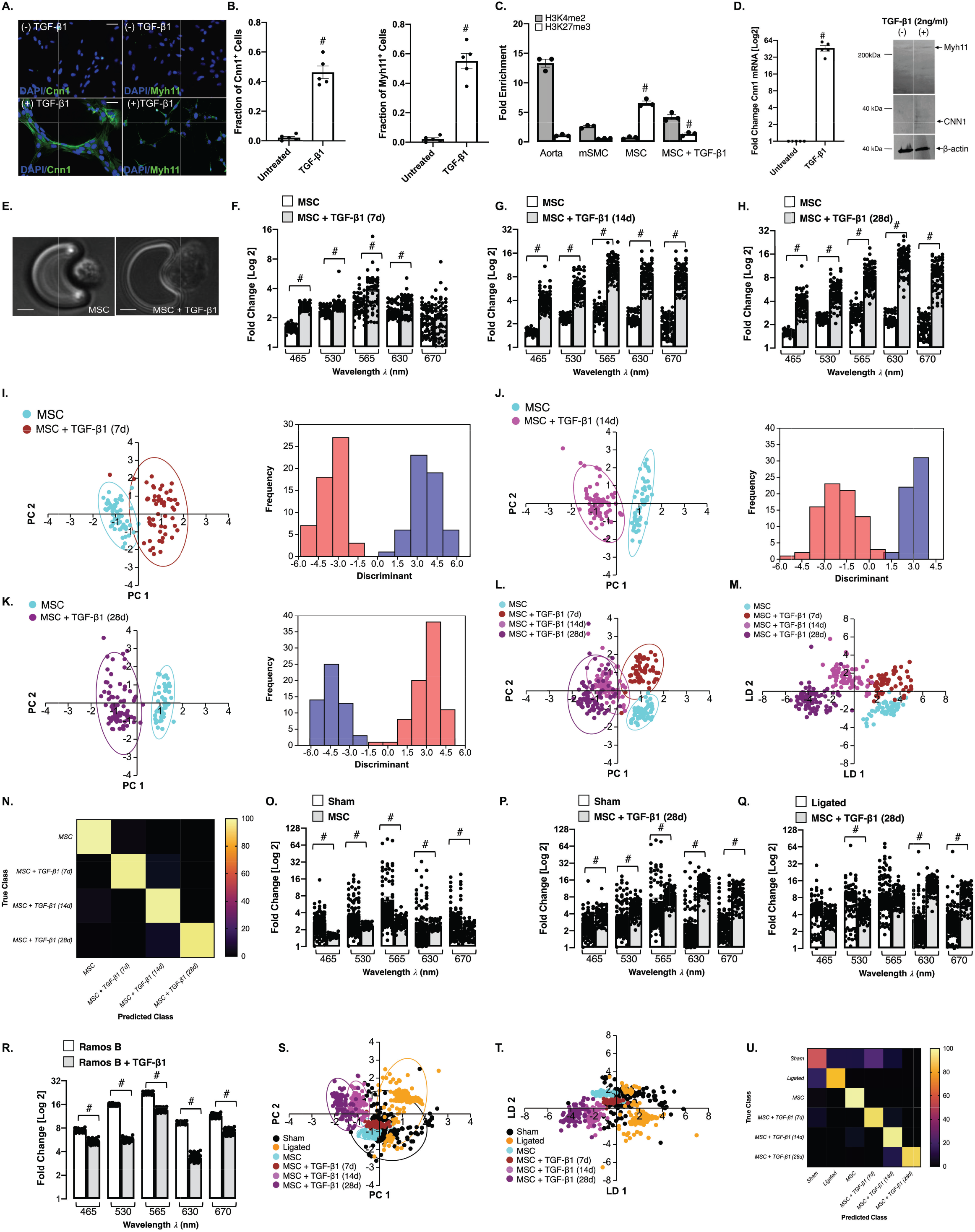
Single cell photonics of murine bone marrow-derived mesenchymal stem cells (mMSCs) before and after myogenic differentiation *in vitro*. **A**. Representative immunocytochemical analysis of Cnn1 (green), Myh11 (green) and DAPI nuclei staining (blue) in mMSCs before and after treatment of cells with media supplemented with TGF-β1 (2ng/mL). B. The fraction of Cnn1^+^ and Myh11^+^ cells before and after treatment of cells with media supplemented with TGF-β1 (2ng/mL). Data are the mean ± SEM of 5 wells, #p<0.05 vs control. C. Fold enrichment of the stable SMC histone modification, H3K4me2 and repressive histone modification, H3K27me3 at the Myh11 promoter in fresh mouse aorta, cultured Movas SMCs and undifferentiated mMSCs in the absence or presence of TGF-β1 (2ng/mL) for 7d. Data are mean ± SEM of a representative experiment performed in triplicate, #p<0.05 vs control (-) TGF-β1 levels. D. Log2 fold change in Cnn1 mRNA levels in mMSCs and representative immunoblot of Cnn1 and Myh11 protein expression in lysates in response to TGF-β1 (2ng/mL). Data are mean ± SEM of n=5, #p≤0.05 considered as significant. E. Visualisation of MSCs and their myogenic progeny on each V-cup in the LoaD platform. **F-H**. Single cell photon emissions from MSCs in the absence or presence of TGF-β1 (2ng/mL) for (F) 7 (G) 14 and (H) 28 d *in vitro*. Data are the Log2 fold increase and represent the mean ± SEM of 55-79 cells/group, #p≤0.001. I-K. LDA plots of MSC (cyan) before and after treatment with TGF-β1 (2ng/mL) for 7d (brown), 14d (magenta) and 28d (purple) cells *in vitro*. Data are the Log2 fold increase and represent the mean ± SEM of 55-79 cells/group, #p<0.001. **L-M**. Combined PCA loading plots and LDA of MSCs and MSCs following treatment with TGF-β1 for 7, 14 and 28d. Data are from 55-79 cells/group. N. Confusion matrix of true class and predicted class following a leave-one-out cross-validation procedure by the LDA classifier. Data are from 268 cells across five wavelengths. O-Q. Single cell photon emissions from MSCs in the absence or presence of TGF-β1 (2ng/mL) 28 d *in vitro* compared to (O,P) sham and (Q) ligated cells *ex vivo*. Data are the Log2 fold increase and represent the mean ± SEM of 55-178 cells/group, #p≤0.001 vs Sham (O,P) Ligated (Q). R. Single cell photon emissions from Ramos B cells in the absence or presence of TGF-β1 (2ng/mL) 14d. Data are the Log2 fold increase and represent the mean ± SEM of 55 cells/group, #p<0.001. **S-T**. PCA loading plots (S) and LDA (T) of sham (black), ligated (orange) and MSC (cyan) before and after myogenic differentiation with TGF-β1 for 7d (brown), 14d (magenta) and 28d (purple). Data are from 624 cells across 5 wavelengths. U. Confusion matrix of true class and predicted class following a leave-one-out cross-validation procedure by the LDA classifier. Data are from 624 cells across 5 wavelengths.

Single cell photonic measurements were recorded in MSCs before and after TGF-β1 treatment for 7, 14 and 28 days, respectively, and visualised by phase contrast microscopy on each V-cup [Figure 3E], Their photonic profiles were compared [Figure 3F-H] and multivariate analysis revealed that MSCs clustered away from their myogenic progeny following differentiation and could be easily separated from their myogenic progeny after 7, 14 and 28 d, respectively [Figure 3I-K] while analysis of the combined data confirmed that MSCs and their myogenic progeny cluster away from each other [Figure 3L] and could be separated over time [Figure 3M] with an accuracy of 96.3% on a cross-validated leave-one-out basis [Figure 3N], When compared with cells isolated from sham and ligated vessels *ex vivo*, the photonic intensity of undifferentiated MSCs was significantly lower than cells from sham vessels [Figure 3O] but increased when these cells were treated with TGF-β1 for 28d [Figure 3P], and was significantly higher at λ630 and λ670 nm when compared to cells from ligated vessels [Figure 3Q]. The increased photonic intensity was not due to the TGF-β1 itself as TGF-β1-treated Ramos B cells had lower photonic intensities across all wavelengths, including at λ565 nm wavelength [Figure 3R]. Multivariate analysis demonstrated a clear separation between MSC-derived myogenic progeny (after 14 and 28d) and cells from ligated vessels [Figure 3S], while undifferentiated MSCs and their myogenic progeny clustered towards cells from sham vessels after 7 d [Figure 3T], Confusion matrices classified 27% of cells from sham vessels and 3% of cells from ligated vessels as similar to MSC-derived myogenic progeny, on a cross-validated leave-one-out basis with 79.1% accuracy [Figure 3U].

Multivariate analysis also revealed that MSCs and their myogenic progeny could be easily separated from Ramos B cells, while MSCs and their myogenic progeny (7d) clustered towards J774A.1 cells [Suppl Figure 2A, B] on a cross-validated leave-one-out basis with 92.8% accuracy [Suppl Figure 2E], consistent with trans-differentiation of macrophages to SMCs in culture [25]. Similarly, MSC-derived myogenic progeny clustered towards aortic SMCs and cultured Movas SMCs [Suppl Figure 2C, D] on a cross-validated leave-one-out basis with 80.1% accuracy [Suppl Figure 2F], Taken together, these data suggest that singlecell photonics clearly discriminates MSCs from their myogenic progeny. Moreover, while SMCs isolated from sham vessels, aortic SMCs, and Movas SMCs share some similarity with MSC-derived myogenic progeny, the photonic profile of the majority of cells isolated from ligated vessels *ex vivo* was distinct.

### Single cell photonics of C3H/10T1/2 mesenchymal stem cells and their myogenic progeny

Myogenic differentiation of C3H/10T1/2 mesenchymal stem cells was confirmed before and after TGF-β1 treatment; SMC differentiation markers Cnn1 and Myh11 increased in response to TGF-β1 [Figure 4A, B] concomitant with a significant increase in Cnn1 and Myh11 expression in protein lysates [Figure 4C] and Cnn1 mRNA levels [Figure 4D],

**Figure 4.**
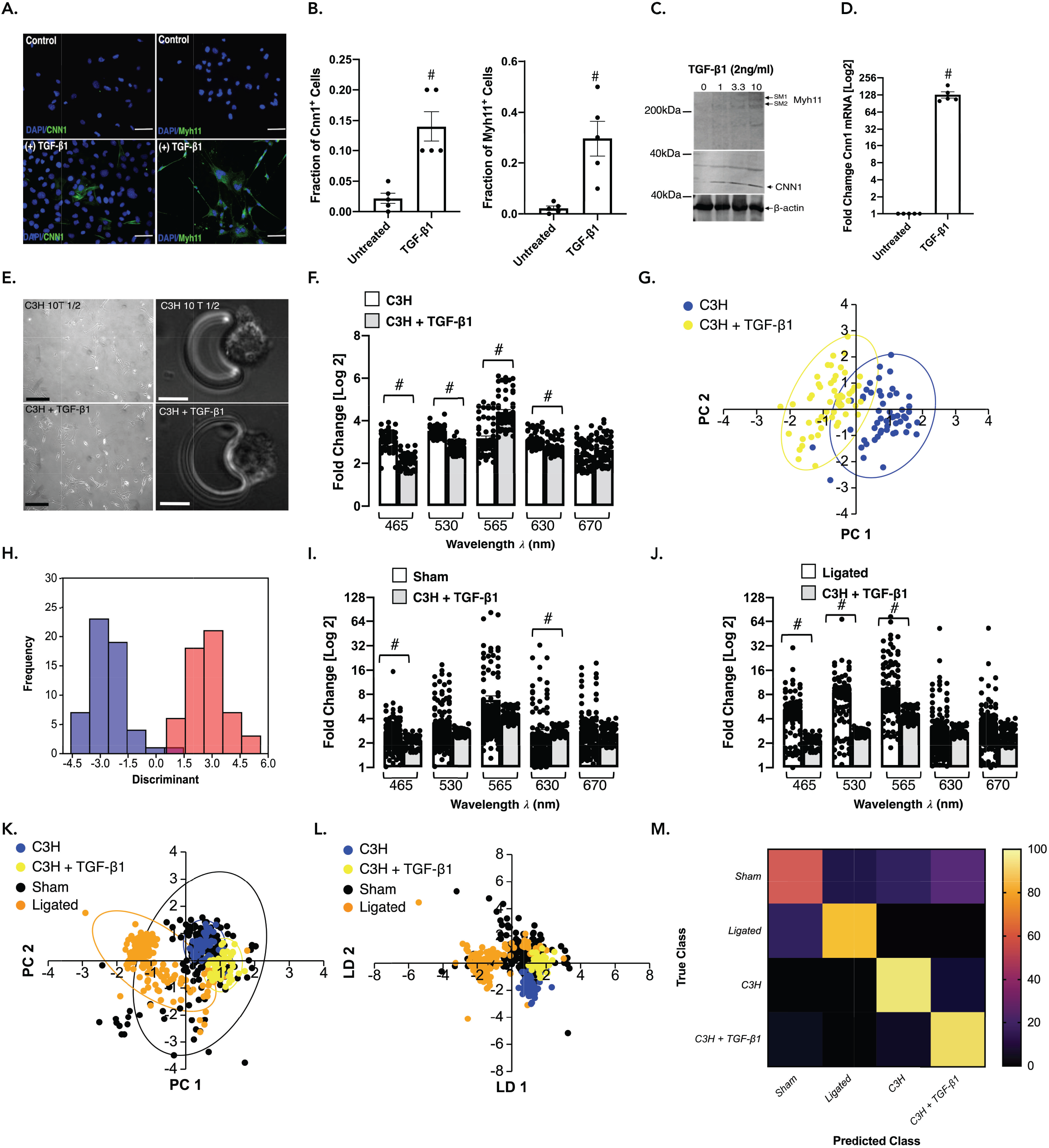
Single cell photonics of C3H/10T1/2 mesenchymal stem cells before and after myogenic differentiation *in vitro*. **A**. Representative immunocytochemical analysis of Cnn1 (green), Myh11 (green) and DAPI nuclei staining (blue) in C3H/10T1/2 cells before and after treatment of cells with media supplemented with TGF-β1 (2ng/mL) for 7d. B. The fraction of Cnn1+ and Myh11^+^ cells before and after treatment of cells with media supplemented with TGF-β1 (2ng/mL). Data are the mean ± SEM of 5 wells, #p≤0.05 vs control. C. Representative immunoblot of Cnn1 and Myh11 protein expression in C3H/10T1/2 lysates in response to TGF-β1 (2ng/mL). D. Log2 fold change in Cnn1 mRNA levels in C3H/10T1/2 cells in response to TGF-β1 (2ng/mL). Data are the mean ± SEM n=5, #p≤0.05. E. Visualisation of C3H10T1/2 cells and their myogenic progeny on each V-cup in the LoaD platform. F. Single cell photon emissions from C3H/10T1/2 in the absence or presence of TGF-β1 (2ng/mL) 7 d *in vitro*. Data are the Log2 fold increase and represent the mean ± SEM of 55 cells/group, #p≤0.001 vs control. **G-H**. PCA loading plots and LDA of C3H/10T1/2 cells before (blue) and after myogenic differentiation with TGF-β1 for 7d (yellow). Data are 55 cells/group. I-J. Single cell photon emissions from C3H/10T1/2 in the absence or presence of TGF-β1 (2ng/mL) 7 d *in vitro* compared to sham (I) and ligated cells (J) *ex vivo*. Data are the Log2 fold increase and represent the mean ± SEM of 55-178 cells/group, #p<0.001. K-L. PCA loading plots and LDA of sham (black), ligated (orange) and C3H/10T1/2 cells before (blue) and after treatment with TGF-β1 (2ng/mL) for 7d (yellow). M. Confusion matrix of true class and predicted class following a leave-one-out cross-validation procedure using the LDA classifier. Data are from 466 cells across five wavelengths.

Single cells were visualised by phase contrast microscopy on each V-cup [Figure 4E] before photonic measurements were recorded before and after myogenic differentiation with TGF-β1 [Figure 4F], Multivariate analysis revealed that undifferentiated C3H/10T1/2 cells clustered away from both their myogenic progeny [Figure 4F] and from Ramos B cells and J774A.1 cells [Suppl Figure 3A, B] and could be separated on a cross-validated leave-one-out basis with 99.3% accuracy.

When compared to cells isolated from sham and ligated vessels [Figure 4I, J], these cells clustered away from ligated cells, but towards sham cells [Figure 4K] and could be separated [Figure 4L] on a cross-validated leave-one-out basis with 75.1% accuracy [Figure 4M].

When compared to aortic SMC *ex vivo* and Movas SMCs, these cells clustered towards aortic SMC and Movas SMCs [Suppl Figure 3C, D] on a cross-validated leave-one-out basis with 78.2% accuracy [Suppl Figure 3G]. In addition, when compared to MSCs and their myogenic progeny, these cells clustered towards MSC-derived myogenic progeny [Suppl Figure 3E, F] on a cross-validated leave-one-out basis with 88.5% accuracy [Suppl Figure 3H].

Taken together, these results suggest that single-cell photonics clearly discriminate C3H/10T1/2 mesenchymal cells from their myogenic progeny. Moreover, while cells isolated from sham vessels share significant similarity with C3H/10T1/2-derived myogenic progeny, the photonic profile of the majority of cells isolated from ligated vessels *ex vivo* were distinct.

### Lineage Tracing analysis of perivascular S100β_+_ vascular stem cells in normal and injured vessels

In order to determine the source of increased medial and intimal cells following ligation injury, lineage tracing analysis was performed using transgenic *S100β-eGFP-CreERT2-Rosa-26-tdTomato* reporter mice [Figure 5A]. The animals were treated with tamoxifen (Tm) for 7 days to induce nuclear translocation of CreER(Tm) and subsequent recombination to indelibly mark resident S100β^+^ mVSc with red fluorescent tdTomato four weeks prior to flow restriction [Figure 5B], Only S100β^+^ mVSc cells present during the period of Tm treatment are marked with tdT and tracked. Morphometric analysis of the *S100β-eGFP-CreERT2-Rosa-26-tdTomato* mice confirmed significant intimal thickening in the ligated left carotid (LCA) compared to the contralateral right carotid artery (RCA) in transgenic mice following flow restriction for 21 days [Figure 5C]. The tissue specificity and recombination efficiency of the Tm-induced Cre activity was confirmed in bone-marrow smears and neuronal tissue from *S100β-eGFP-CreERT2-Rosa-26-tdTomato* mice [data not shown].

**Figure 5.**
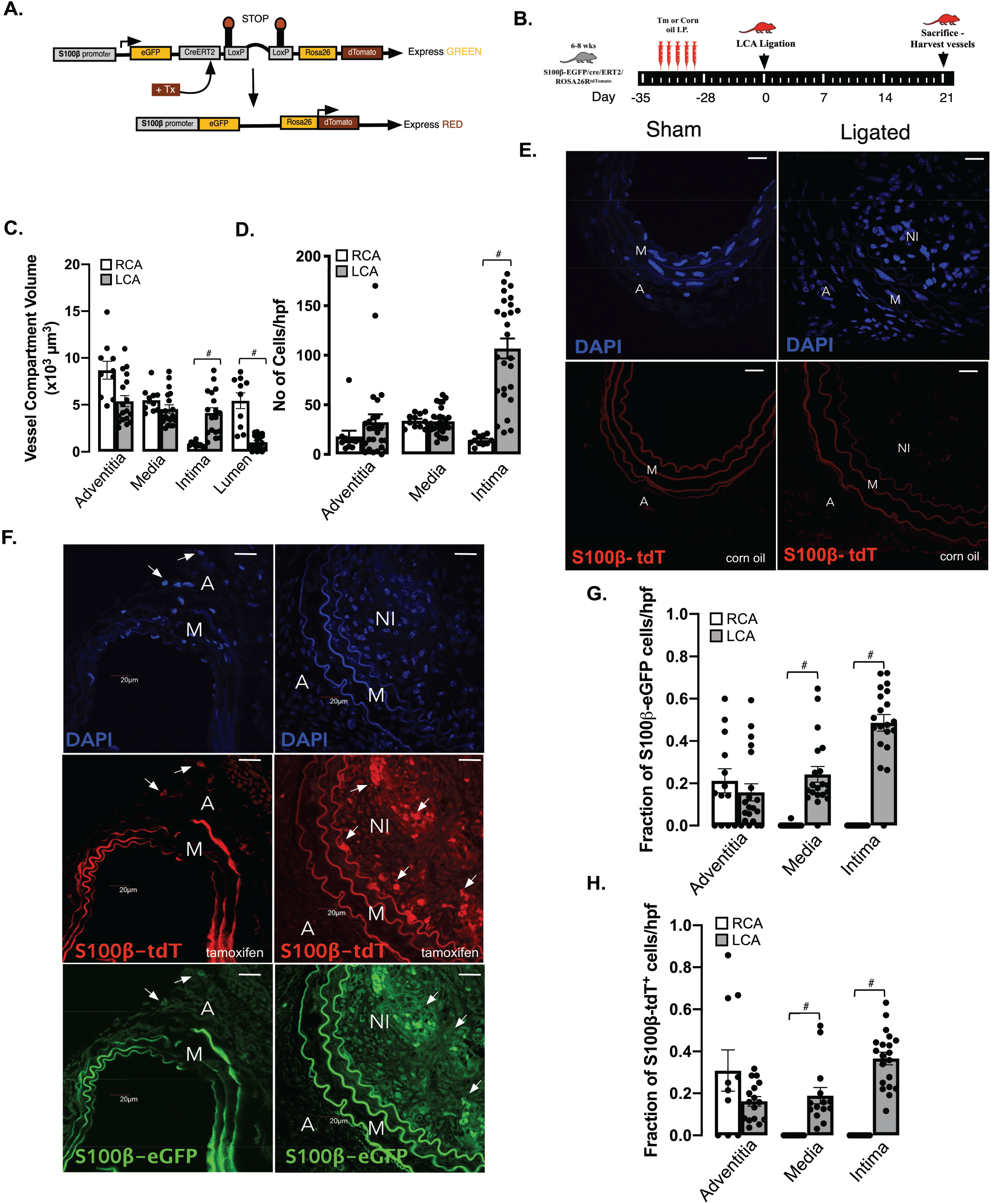
Lineage tracing analysis of marked perivascular S100β cells following flow restriction. **A**. Schematic diagram showing genetic lineage tracing using S100β-CreERT2-Rosa26-tdTomato. **B**. Schematic diagram showing protocol for Tm injections before ligation after 4 week Tm washout. **C**. Morphometric analysis of adventitial, medial, intimal and luminal volumes in male and female S100β-CreERT2-Rosa26-tdTomato mice following complete ligation of the left carotid artery (LCA) for 21 d. Data are from 4 animals per experimental group. D. Representative confocal fluorescence images of DAPI nuclei (blue) and S100β-tdT^+^ (red) cells in sham and ligated LCA in corn oil (control) treated animals. Representative images shown; 4 animals per experimental group. Scale bar = 50μm. **E**. Representative confocal fluorescence images of DAPI nuclei (blue), S100β-tdTΓ (red) cells S100β-eGFP^+^ cells in sham and ligated LCA in tamoxifen (Tm) treated animals. Representative images are from 4 animals per experimental group. Scale bar = 50μm. **F**. Cumulative analysis of the number of DAPI nuclei/hpf in the adventitial, medial and intimal layers from the contralateral RCA and ligated LCA. Data are the mean ± SEM of 10-20 images/group, #p≤0.05. **G**. Cumulative analysis of the fraction of S100β-eGFP^+^ cells within the adventitia (A), media (M), and neointima (NI) of RCA and ligated LCA. Data are the mean ± SEM of 15-20 sections per experimental group, #p≤ 0.05 from 4 animals per group. **H**. Cumulative analysis of the fraction of S100β-tdT^+^ cells within the adventitia (A), media (M), and neointima (NI) of RCA and ligated LCA. Data are the mean ± SEM of 15-20 sections per experimental group, #p≤ 0.05 from 4 animals per group.

Treatment of *S100β-eGFP-creER2-Rosa26tdT* transgenic mice with Tm indelibly marked S100β^+^ cells (S100β-tdT^+^) within the adventitial layer of the LCA prior to flow restriction [Figure 5D], No cells are marked when these transgenic mice were treated with the vehicle control (corn oil) [Figure 5D], Importantly, no S100β-tdT^+^ cells were observed in the intimal (EC) or medial (SMC) layers of vessels following tamoxifen treatment before flow restriction [Figure 5D], However, following ligation, there was a striking increase in the number of S100β-tdT marked cells within the medial and intimal layers of the ligated LCA, compared to RCA control [Figure 5E], Cumulative analysis confirmed a significant increase in the number [Figure 5F] and the fraction of S100β-eGFP^+^ cells within the LCA medial and intimal layers, compared to the RCA control [Figure 5G]. Moreover, a significant proportion (~40%) of these cells originated from an S100β-tdT marked parent population since they were also marked with tdT [Figure 5H], Since intimal and medial cells are not marked with S100β-CreER tdT prior to injury, these data suggest that a significant proportion of intimal and medial cells are derived from a S100β^+^ (non-SMC) parent population following flow restriction *in vivo*.

### Single cell photonics of S100β_+_ resident vascular stem cells and their myogenic progeny

Resident S100β^+^ mVSc were isolated from murine aorta by sequential plating. Immunocytochemical analysis confirmed that these cells were S100β positive [Suppl Figure 4A], Cnn1 protein [Figure 6A, B], Cnn 1 mRNA expression [Figure 6C] and enrichment of stable SMC epigenetic H3K4me2 histone mark at the Myh11 promoter [Figure 6D] significantly increased after treatment for 7 d with media supplemented with either TGF-β1 (2ng/mL), Jag1 (1.0μg/mL), or SHh (0.5μg/mL).

**Figure 6.**
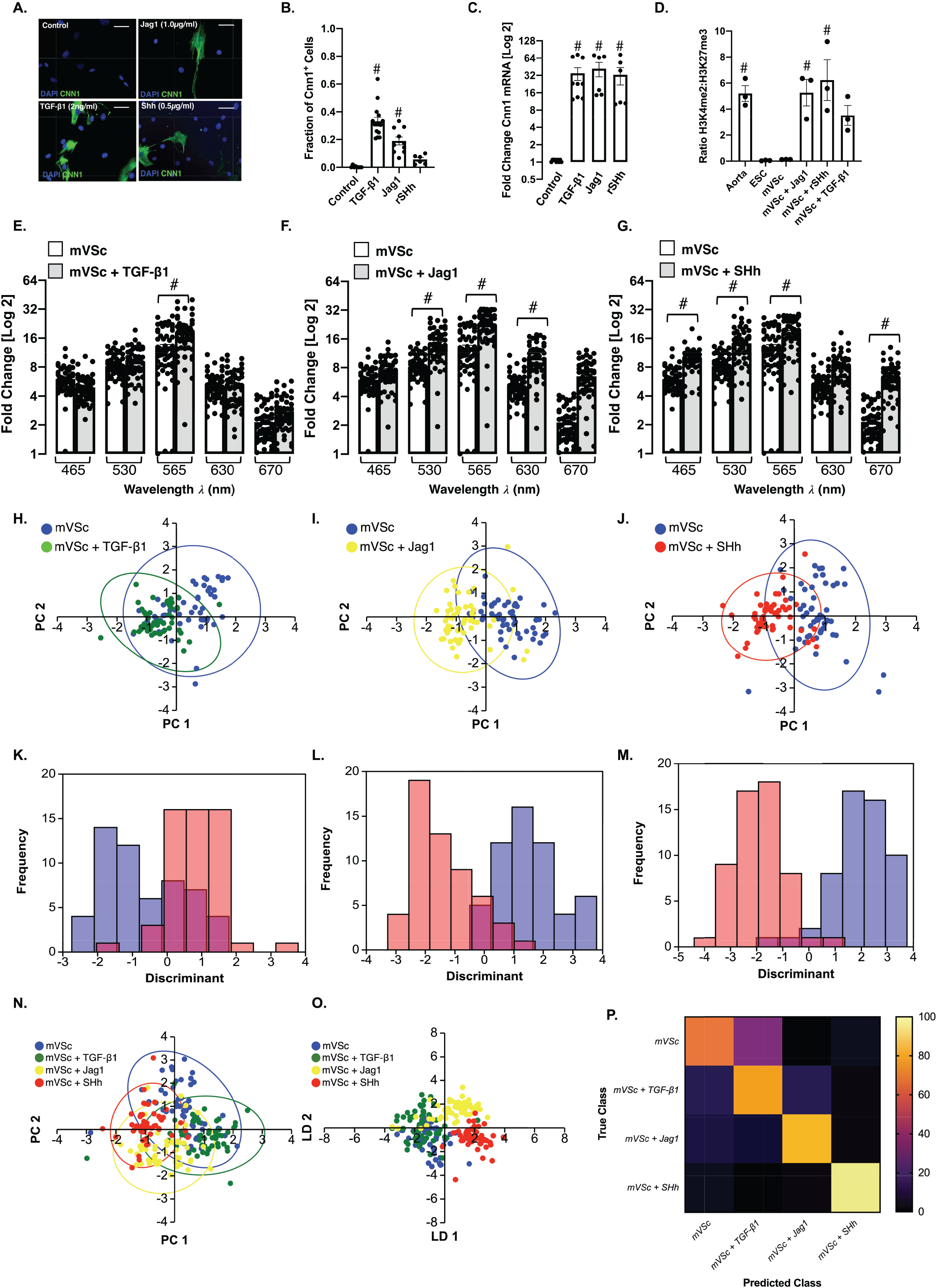
Single cell photonics of S100β^+^ vascular stem cells (mVSc) before and after myogenic differentiation *in vitro*. **A**. Representative immunocytochemical analysis of Cnn1 (green) and DAPI nuclei staining (blue) in mVSc before and after treatment of cells with media supplemented with TGF-β1 (2ng/mL), Jag1 (1μg/mL) and SHh (0.5μg/mL) for 14d. **B**. The fraction of Cnn1^+^ cells before and after treatment of cells with media supplemented with TGF-β1 (2ng/mL), Jag1 (1μg/mL) and SHh (0.5μg/mL) for 14d. Data are the mean ± SEM of 3 experiments, #p≤0.05 vs control. **C**. Log2 fold change in Cnn1 mRNA levels before and after treatment of cells with media supplemented with TGF-β1 (2ng/mL), Jag1 (1μg/mL) and SHh (0.5μg/mL) for 7d. Data are the mean ± SEM n=3, #p≤0.05 vs control. **D**. Fold enrichment of the stable SMC histone modification, H3K4me2 and repressive histone modification, H3K27me3 at the Myh11 promoter in fresh mouse aorta (Aorta), mouse embryonic stem cells (ESC), and undifferentiated S100β^+^ mVSc before (mVSc) and after treatment of cells with media supplemented with TGF-β1 (2ng/mL), Jag1 (1μg/mL) and SHh (0.5μg/mL) for 7 d. Data are representative of n=3, #p<0.05 vs control. E-G. Single cell autofluorescence photon emissions from mVSc in the absence or presence TGF-β1 (2ng/mL), Jag1 (1μg/mL) and SHh (0.5μg/mL) for 14d *in vitro*. Data are the Log2 fold increase and represent the mean ± SEM of 55 cells/group, #p≤0.001 vs control. H-J. Individual PCA loading plots of mVSc before (blue) and after myogenic differentiation with (H) TGF-β1 (green), (I) Jag1 (yellow) and (J) SHh (red). K-M. Individual LDA plots of mVSc before and after myogenic differentiation with (K) TGF-β1, (L) Jag1 and (M) SHh. N. Cumulative PCA loading plots of mVSc before (blue) and after myogenic differentiation with (H) TGF-β1 (green), (I) Jag1 (yellow) and (J) SHh (red). O. Cumulative LDA plots of mVSc before and after myogenic differentiation with (K) TGF-β1, (L) Jag1 and (M) SHh. P. Confusion matrix of true class and predicted class following a leave-one-out cross-validation procedure by the LDA classifier. Data are from 219 cells across five wavelengths.

The photonic intensity of single cells captured on each V-cup [Suppl Figure 4B] at the λ565 nm wavelength following myogenic differentiation for 14 d increased in response to all three myogenic inducers [Figure 6E-G]. Multivariate analysis revealed that S100β^+^ mVSc clustered away from their respective myogenic progeny following differentiation [Figure 6H-J] and could be separated on a cross-validated leave-one-out basis (TGF-β1:77.1%, Jag-1: 90% and SHh: 93.4% accuracy) [Figure 6K-M], Multivariate analysis of the combined data confirmed S100β^+^ mVSc could be easily discriminated from their myogenic progeny [Figure 6N,O] when cross-validated on a leave-one-out basis with 79.5% accuracy [Figure 6P].

When compared to aortic SMC and Movas SMCs, S100β^+^ mVSc and their myogenic progeny clustered away from these cells [Suppl Figure 4C] and could be easily separated [Suppl Figure 4D] on a cross-validated leave-one-out basis with 76.3% accuracy [Suppl Figure 4E], Similarly, multivariate analysis confirmed that S100β^+^ mVSc and their myogenic progeny could be discriminated from J774A.1 and Ramos B cells [Suppl Figure 4F-H] and MSCs and their myogenic progeny [Suppl Figure 5A-C], on a cross-validated leave-one-out basis with 85.7% and 88.3% accuracy, respectively.

When compared to cells isolated from sham [Figure 7A-C] and ligated vessels [Figure 7D-F], multivariate analysis of the spectra revealed that S100β^+^ mVSc and their myogenic progeny clustered away from sham cells but towards isolated cells of ligated vessels [Figure 7G, H] and could be discriminated on a cross-validated leave-one-out basis with 73.9% accuracy [Figure 7I-K],

**Figure 7.**
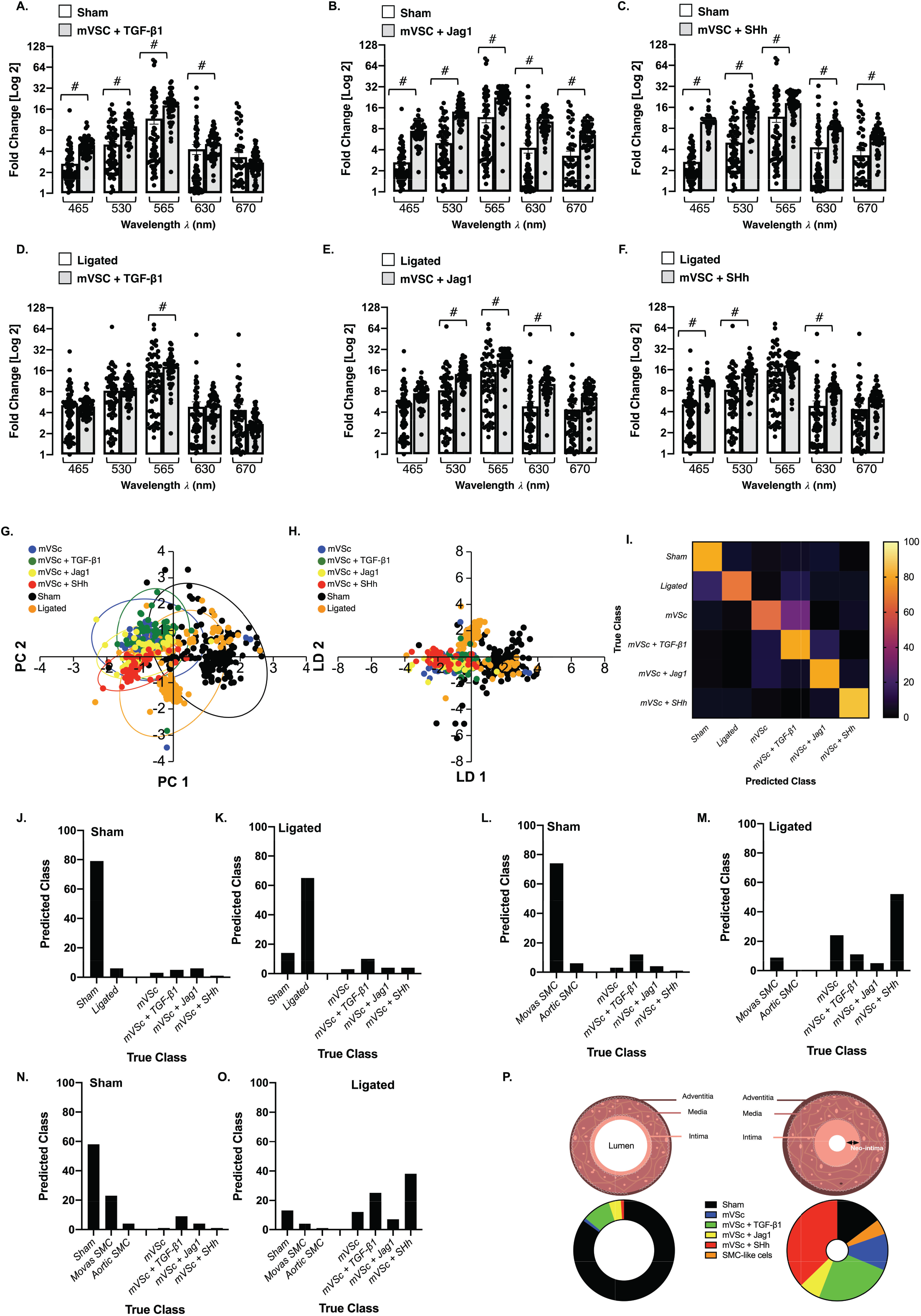
Single cell photonics identifies S100β resident vascular stem cells and their myogenic progeny within ligated vessels. **A-C**. Single cell photon emissions from Sham cells *ex vivo* compared to mVSc in the presence (A) TGF-β1 (2ng/mL), (B) Jag1 (1μg/mL) and (C) SHh (0.5μg/mL) for 14 d *in vitro*. Data are the Log2 fold increase and represent the mean ± SEM of 55-178 cells/group, #p≤0.001 vs control. D-F. Single cell auto-fluorescence photon emissions from ligated cells *ex vivo* compared to mVSc in the presence (D) TGF-β1 (2ng/mL), (E) Jag1 (1μg/mL) and (F) SHh (0.5μg/mL) for 14 d *in vitro*. Data are the Log2 fold increase and represent the mean ± SEM of 55-178 cells/group, #p≤0.001 vs control. **G-H**. Cumulative PCA loading plots and LDA of sham (black), ligated (orange) and mVSc before (blue) and after myogenic differentiation with (H) TGF-β1 (green), (I) Jag1 (yellow) and (J) SHh (red). I. Confusion matrix of true class and predicted class following a leave-one-out cross-validation procedure by the LDA classifier. Data are from 575 cells across five wavelengths. J-K. Percentage of cells in sham and ligated vessels predicted as mVSc and their myogenic progeny by LDA classifier. Data are from 575 cells across five wavelengths **L-M**. Percentage of cells in sham and ligated vessels classified as aortic SMCs, Movas SMCs and mVSc and their myogenic progeny when trained with datasets by LDA classifier. Data are from 356 cells. N-O. Percentage of cells in sham and ligated vessels classified as sham cells, aortic SMCs, Movas SMCs and mVSc and their myogenic progeny by the LDA classifier. Data are from 356 cells across five wavelengths. P. Graphic representation of percentage of cells in sham and ligated vessels by LDA classifier.

We also analysed cell diameter and evaluated whether a correlation exits between cell shape changes and autofluorescence (AF) emissions. The cell diameter of medial cells from sham vessels (both carotid artery and aorta) was similar but significantly lower compared to cells from ligated vessels (Suppl Figure 6). While cell diameter increased following myogenic differentiation in response to TGF-β1, an effect not mirrored in Ramos B cells following similar treatment, the changes in cell diameter were not significantly correlated with autofluorescence emissions at λ565 nm (data not shown).

### Single cell photonics identifies S100β_+_ resident vascular stem cells and their myogenic progeny within ligated vessels

In order to assess the contribution of S100β^+^ mVSc and their myogenic progeny to the overall heterogenous phenotype of cells isolated from sham and ligated vessels, the photonic profiles of (i) Movas SMCs, (ii) aortic SMCs and (iii) S100β^+^ mVSc and their myogenic progeny were used as part of a training set before sham and ligated vessel cells were interrogated against this trained dataset using LDA. LDA of spectra from 685 cells separated each group on a cross-validated leave-one-out basis with 66.7 % accuracy. In sham vessels, the majority of cells were classified as Movas SMC [Figure 7L]. In contrast, the majority of cells within ligated vessels were classified as S100β^+^ mVSc or their myogenic progeny (~65%) with the remainder classified as Movas SMC [Figure 7M], When cells isolated from sham vessels were also included in this training set, the majority of sham cells were classified as SMC (either sham, Movas or aortic SMCs) [Figure 7N], In contrast, in ligated vessels, the majority of cells were classified as S100β^+^ mVSc and their myogenic progeny on a cross-validated leave-one-out basis with 66.6% accuracy [Figure 7O]. Taken together, these data suggest that a significant proportion of cells within ligated/remodelled vessels share a photonic profile similar to S100β^+^ mVSc and their myogenic progeny [Figure 7P].

When the contribution of all cell types was analysed by LDA as part of an overall training set before single cells isolated from ligated vessels were interrogated, LDA of spectra from 1063 cells separated each group on a cross-validated leave-one-out basis with 67.7% accuracy [Figure 8A], The majority of cells isolated from sham vessels were classified as themselves or Movas SMC, MSCs or their TGF-β1-derived myogenic progeny and compromised ~86% of all cells [Figure 8B], When individual cells isolated from ligated vessels were interrogated against this trained dataset, a significant number of ligated cells were classified as S100β^+^ mVSc and their TGF-β1-, Jag1- and SHh-derived myogenic progeny [Figure 8C].

**Figure 8.**
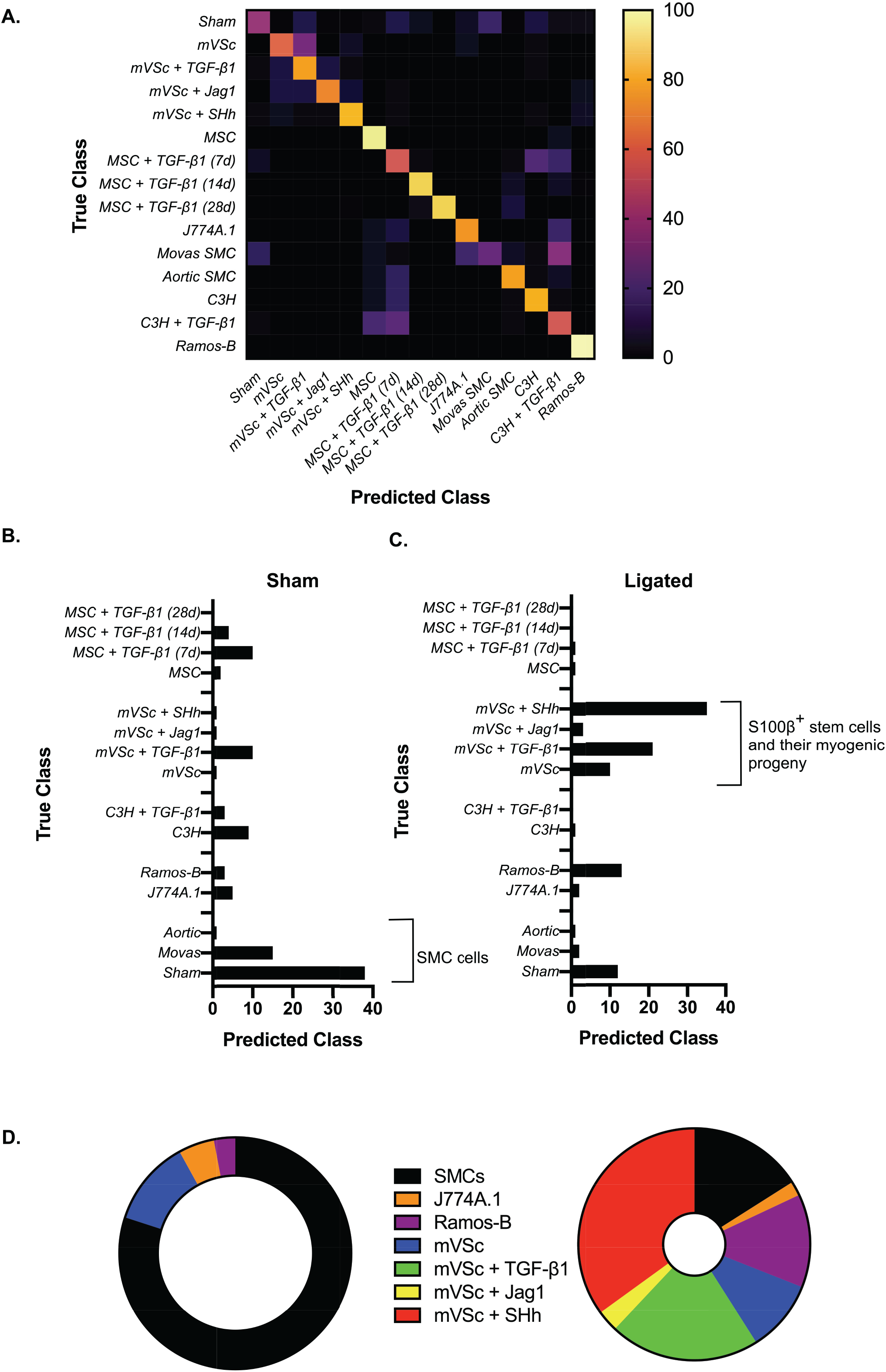
Single cell photonics of all cells identifies S100β resident vascular stem cells and their myogenic progeny within ligated vessels using an LDA classifier. **A**. Confusion matrix of true class and predicted class of all cells following a leave-one-out cross-validation procedure by the LDA classifier. The LDA algorithm was trained using single cell photonic analysis from (i) sham cells, (ii) aortic SMCs, (iii) Movas SMCs, (iv) J774A.1 macrophages, (v) Ramos B cells, (vi) MSCs and their myogenic progeny, (vii) S100β mVSc and their myogenic progeny. Data are from 885 cells B. Percentage of cells in sham vessels using the LDA classifier when trained with all cells. Data are from 178 cells. C. Percentage of cells in ligated vessels using the LDA classifier when trained with all cells. Data are from 178 cells **D**. Graphic representation of percentage of cells in sham and ligated vessels when trained with all cells using the LDA classifier.

These data suggest that single-cell photonic profiles *ex vivo* are of sufficient coverage, specificity and overall quality for LDA algorithms to discriminate various different cell types within injured vessels *ex vivo* and reveal that a significant number of ligated cells were classified as S100β^+^ mVSc and their myogenic progeny *in vitro* [Figure 8D].

### Supervised machine learning to interrogate isolated cells from normal and injured vessels

As part of a supervised machine learning procedure, MLP artificial neural network analysis was subsequently deployed to further classify each cell population and interrogate cells from sham and ligated vessels [26] [Figure 9A], To train the network, we optimized several experimental conditions, including number of wavelengths for input data, number of hidden layers, the momentum and learning rate. The final classification model had an F-score of 0.928, recall of 0.929 and precision score of 0.929 [Figure 9B], Cross validation confirmed that optimized neural networks can identify all cell types with high performance, based only on their autofluorescence emissions and resulted in an F-score of 0.81. The cells were also trained on a 66 % split before the remainder was tested and the F score dropped to 0.77. This trained data set was subsequently used to interrogate spectral datasets of cells isolated from sham and ligated vessels *ex vivo*. The majority of sham cells (>80%) were classified as sham [Figure 9C]. In contrast, ~50% of ligated cells were classified as mVSc and their SHh-derived myogenic progeny in addition to sham cells (~19%) and non-classified ligated cells [Figure 9D, E].

**Figure 9.**
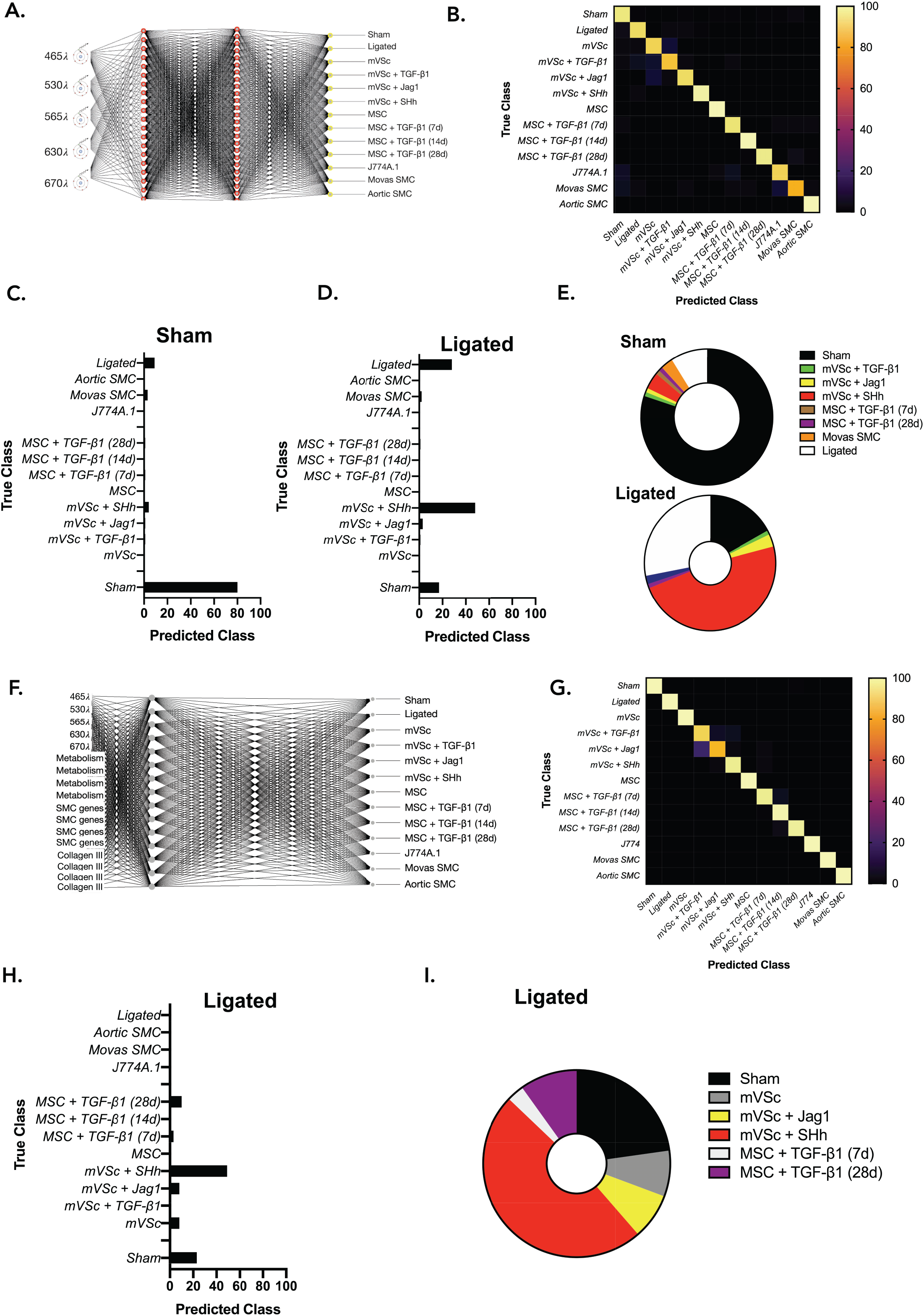
Supervised machine learning identifies S100β^+^ mVSc-derived myogenic progeny within ligated vessels. **A**. Graphic representation of the MLP neural network algorithm. **B**. Confusion matrix of true class and predicted class of following training with (i) sham cells, (ii) ligated cells (iii) aortic SMCs, (iv) Movas SMCs, (v) J774A.1 macrophages, (vi) MSCs and their myogenic progeny and (vii) S100β mVSc and their myogenic progeny using MLP neural network analysis. **C-D**. Percentage of cells in sham and ligated vessels classified using MLP neural network analysis on trained dataset. The trained dataset consisted of 924 cells across five wavelengths. The test dataset consisted of 78 cells across five wavelengths E. Graphic representation of percentage of cells in sham and ligated vessels using MLP neural network analysis. F. Graphic representation of the MLP neural network algorithm combining photonic signatures with indices of cell metabolism, lineage specific gene expression and structural gene expression to predict the cellular heterogeneity within vascular lesions. **G**. Confusion matrix of true class and predicted class of following training with (i) sham cells, (ii) ligated cells (iii) aortic SMCs, (iv) Movas SMCs, (v) J774A.1 macrophages, (vi) MSCs and their myogenic progeny and (vii) S100β mVSc and their myogenic progeny using re-trained dMLP neural network analysis. H. Percentage of cells in ligated vessels classified using MLP neural network analysis on re-trained dataset. The trained dataset consisted of 924 cells across five wavelengths. The test dataset consisted of 78 cells across five wavelengths I. Graphic representation of percentage of cells in ligated vessels using MLP neural network analysis.

As undifferentiated cells can switch between mitochondrial respiration and aerobic glycolysis (even in the presence of oxygen) akin to the “Warburg effect” whereby cells increase their lactate output to fuel their ever-increasing metabolic demands, we determined the metabolic state of undifferentiated stem cells before and after treatment with a myogenic stimulus *in vitro*. The levels of glucose and glutamine consumption in addition to lactate release relative to protein content were all evaluated in mVSc conditioned media before and after myogenic differentiation with the Notch ligand, Jag-1. Myogenic differentiation was associated with a significant increase in glucose consumption and enhanced lactate release over time when compared to undifferentiated cells up to day 14. In contrast, glutamine consumption was similar until day 14 (Suppl Figure 7A-C). This was specific for myogenic differentiation as inhibition of Jag-1 induced Notch signalling with DAPT inhibited these glycolytic responses (Suppl Figure 7D-F).

Our new network was then re-trained to include the level of glycolytic metabolism, SMC differentiation marker gene and Col 3A1 expression as additional variables to the photonic autofluorescence emissions of cells as a predictor of phenotype within lesions. To re-train the network, we again optimized several all experimental conditions. The number of hidden layers was optimised at 1 and the final re-trained classification model had an improved F-score of 0.997, recall of 0.997 and precision score of 0.987. Cross validation confirmed that the optimized neural network can identify all cell types with high performance, based on their autofluorescence emissions, SMC differentiation gene expression, Col 3A1 levels and glycolytic metabolism and resulted in an F-score of 0.975. The cells were also trained on a 66 % split before the remainder was tested and the F-score was improved at 0.981 (Figure 9F-G). This re-trained data set was subsequently used to interrogate single cells isolated from ligated vessels *ex vivo*. Again, the majority of cells from ligated vessels were classified as mVSc and their SHh-derived myogenic progeny (Figure 9H, I).

Thus, supervised machine learning using MLPs neural network analysis successfully identified each cell type in both training sets with a high degree of accuracy (>90%) and further confirmed lineage tracing analysis that a significant number of ligated cells *ex vivo* are classified as S100β^+^ mVSc and their myogenic progeny *in vitro* [Figure 9E, I].

### Elastin and Coll 3A1 contribute to the photonic intensity of stem cell-derived myogenic progeny at λ565 ± 20 nm wavelength

PCA loading plots of cells isolated from sham and ligated vessels suggest that the photonic differences at λ565 ± 20nm had a significant influence on PC1 [Figure 1I]. Similarly, the photonic profile at λ565 ± 20nm had a significant influence on PC1 in PCA biplots of single MSCs, S100β^+^ mVSc and C3H/10T1/2 cells following myogenic differentiation *in vitro* when compared [Figure 10A], When the photonics of single cells isolated from sham and ligated vessels was compared to MSCs, S100β^+^ mVSc and C3H/10T1/2 cells at λ565 ± 20nm, there was a marked increase in the intensity at this wavelength in cells isolated from ligated vessels and cells undergoing myogenic differentiation [Figure 10B]. In order to identify one of the fluorophores that may contribute to the increase in the photonic intensity of stem cell-derived myogenic progeny at λ565 ± 20 nm, the roles of two important auto fluorescent molecules, elastin and Coll3A1 were assessed [27], The mRNA levels of elastin and Coll3A1 were both significantly increased following myogenic differentiation of mMSCs with TGF-β1 treatment [Figure 10C] and S100β^+^ mVSc with TGF-β1 [Figure 10D] and Jag1 [Figure 10E]. The effects of elastin and Coll3A1 depletion on the photonic profile of undifferentiated mMSCs at the λ565 ± 20 nm wavelength was also assessed following transfection of cells with specific siRNA duplexes targeting elastin and Coll3A1 transcripts, respectively. A fluorescently tagged siRNA was used to validate transfection efficiency at over 80% [data not shown] before elastin and Coll3A1 mRNA depletion following TGF-β1 treatment was confirmed by qRT-PCR after 72 h [Figure 10F].

**Figure 10.**
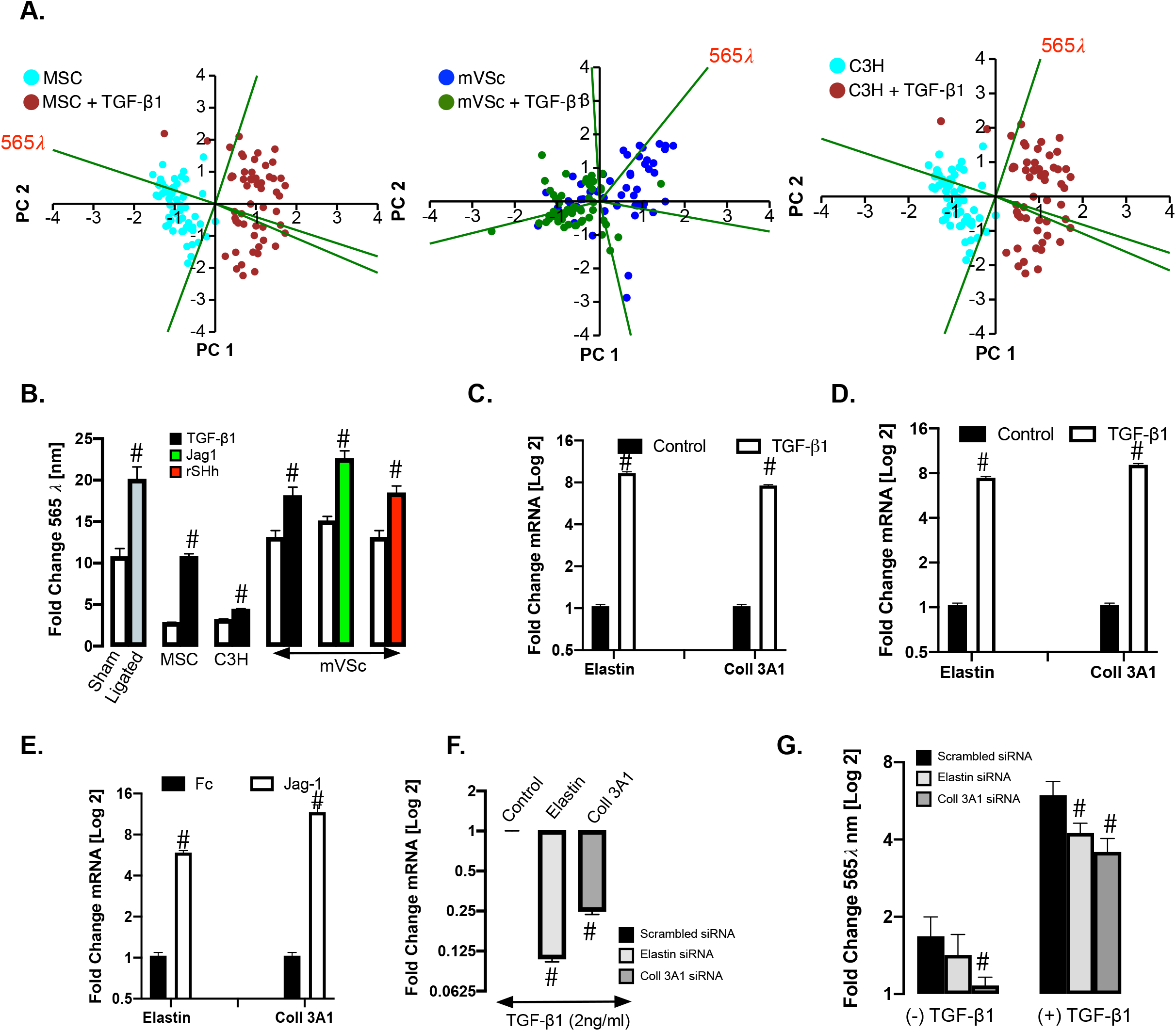
Elastin and Coll 3A1 contribute to the photonic intensity of stem cell-derived myogenic progeny at λ565 ± 20 nm wavelength. **A**. PCA biplots of single cell photon analysis from MSC, mVSc and C3H/10T1/2 cells in the absence or presence of TGF-β1. Each *λ* variable in the PCA biplot is representative of the eigenvectors (loadings). The λ565 variable strongly influences PC1 when cells undergo myogenic differentiation. B. Log2 fold changes in single cell photon emissions at λ565 nm in sham and ligated cells *ex vivo*, MSC and mVSc following myogenic differentiation. Data are the mean ± SEM of 55-178 cells/group, #p≤0.01. C. Log2 fold changes in mRNA levels for elastin and Coll3A1 in MSCs following treatment with TGF-β1 for 7 d. Data are the mean ± SEM of n= 3, #p≤0.05 vs control. D. Log2 fold changes in mRNA levels for elastin and Coll 3A1 in mVSc following treatment with TGF-β1 for 7 d. Data are the mean ± SEM of n= 3, #p≤0.05 vs control E. Log2 fold changes in mRNA levels for elastin and Coll 3A1 in mVSc following treatment with Jag1 for 7 d. Data are the mean ± SEM of n= 3, #p≤0.05 vs Fc control. F. Log2 fold changes in mRNA levels for elastin and Coll3A1 in response to TGF-β1 (2ng/mL) following siRNA knockdown of elastin and Coll3A1, respectively. Data are the mean ± SEM of n= 3, #p≤0.05 vs scrambled control. G. Log2 fold changes in single cell photon emissions at λ565 nm in response to TGF-β1 (2ng/mL) following siRNA knockdown of elastin and Coll3A1, respectively. Data are the mean ± SEM of n= 3, #p≤0.05 vs scrambled control.

In order to establish whether collagens might contribute to the photonic intensity of myogenic progeny at λ565 ± 20nm, we performed spectral analysis using the LoaD platform on various forms of recombinant collagen (Col |⍰1, Col |⍰2 and Col 3A1) as these collagens are synonymous with vascular tissue [28]. In particular, Col I and 3A1 are enhanced in SMCs in culture and within vascular lesions [29, 30]. While autofluorescence emissions were detected for all collagens examined, the predominant autofluorescence emissions at λ560 ± 20 nm were detected for Col 3A1 (Suppl Figure 8A). To further address these collagen spectra more efficiently, spectra across various excitatory wavelengths (λ400-700 nm) were analysed following removal of background emissions due to PBS. Peak Col 3A1 autofluorescence emissions were predominantly observed between λ400 – λ480⍰nm and λ510 – λ570 nm using λ358⍰ and λ 488nm excitation, respectively (Suppl Figure 8B-F). When Col 3A1 was excited at various wavelengths (λ300-500 nm), clear emission spectra were observed between at λ465, λ530 and λ565 nm (Suppl Figure 8G-I). Notably, the photonic intensity of undifferentiated cells at λ565 ± 20nm following Coll3A1 depletion was significantly decreased without a significant effect following elastin depletion, when compared to the scrambled siRNA controls [Figure 10G]. However, the photonic intensity at λ565 ± 20 nm was significantly reduced following treatment with TGF-β1 for 7 d in both elastin and Coll3A1 depleted mMSCs [Figure 10G].

These data confirm that the autofluorescence emissions at λ565 ± 20nm during myogenic differentiation of undifferentiated stem cells was due, in part, to changes in elastin an Coll3A1 expression within these cells.

### Discussion

The application of single cell autofluorescence to study vascular physiology and pathology is an exciting emerging field [31, 32], The ability to easily discriminate heterogeneous populations of cells in the context of various disease processes is of potential clinical and diagnostic value. Herein, we demonstrate the feasibility of multivariate analysis of single cell photonics as a powerful discriminator of cell phenotype using optical multi-parameter interrogation of single cells on a novel Lab-on-a-Disk (LoaD) platform. Single cell photonics were of sufficient coverage, specificity, and quality to discriminate various disparate cell phenotypes *in vitro* and normal medial SMCs from SMC-like cells following injury *ex vivo*, in addition to distinguishing myogenic differentiation of a series of multipotent stem cells *in vitro*. Using these photonic datasets, cellular heterogeneity within vascular lesions was confirmed by identifying the presence of S100β^+^ stem-derived myogenic progeny using multivariate analysis and supervised machine learning and validated by genetic lineage tracing *in vivo*. A combination with indices of cell metabolism, lineage specific gene expression and structural gene expression further enhanced the AI model in predicting the cellular heterogeneity within vascular lesions.

In this study, we clearly demonstrate the feasibility of measuring specific autofluorescence emissions from single cells to discriminate several cell phenotypes *ex vivo* or *in vitro*. In particular, differentiated medial SMCs displayed significant differences in their photonic profile across five broadband light wavelengths when compared to SMC-like cells from injured vessels, SMCs in culture, undifferentiated stem cells, macrophages and B-cells. Notably, significant differences were observed in medial SMCs from normal aortic and carotid vessels. Based on these data, it is clear that considerable heterogeneity exists in the photonic profile of medial SMCs between vascular beds in healthy animals, and yet medial SMCs uniformly express SMC contractile markers (SMA and Myh11). This heterogeneity is noteworthy as it may indicate the existence of specific subsets of medial SMCs with particular disease-relevant photonic profiles. Similar conclusions were observed using scRNA-seq data as a discriminator [33]. As carotid artery SMCs are atheroprone and are derived from the neural crest [34] whereas medial SMCs derived from the thoracic aortic are of somatic mesoderm origin [35] and are considered athero-resistant, it is likely that differences in their embryological origin may contribute to these distinct photonic profiles.

Bone-marrow derived MSCs and C3H/10T1/2 cells are considered good models of SMC differentiation from mesoderm precursors *in vitro* [39, 40] and single cell autofluorescence emissions clearly discriminated undifferentiated stem cells from their myogenic progeny. These differences were also apparent when myogenic progeny were interrogated by Raman and Fourier Transform Infrared (FTIR) spectroscopy as a discriminator where they shared a similar photonic profile with ‘de-differentiated’ sub-cultured vascular SMCs *in vitro* [38]. It is also not surprising that MSC- and C3H/10T1/2-derived myogenic progeny shared similar photonic characteristics to aortic SMCs *ex vivo* and cultured Movas SMCs since they are derived from the same mesoderm origin. As resident S100β mVSc are neuroectoderm in origin and express neural stem cell markers such as S100β, Nestin, So×10 and So×17 [39–41], these cells and their myogenic progeny were distinct to aortic SMCs based on their photonic profile. Furthermore, it is unlikely these resident stem cells are derived from medial SMCs as they did not enrich for the stable SMC epigenetic histone mark, H3K4me2 at the Myh11 locus [23]. However, these myogenic progeny were most notable for their photonic similarity to cells of ligated remodelled vessels.

For many years, SMC-like cells within neointimal lesions have been considered monoclonal/oligoclonal in origin arising from the expansion of a pre-existing resident clonal population of cells. Despite extensive research, their origin remains controversial. Although the Myh11-CreERT2 transgene was originally deemed specific for vascular SMC cell fate mapping studies [42], data indicating expression of this and other SMC differentiation genes in non-SMC populations has recently emerged [43–45] raising the possibility that other resident cells may contribute to lesion formation. In this study, we provide compelling genetic evidence using lineage tracing analysis that vascular lesions contain an abundance of S100β^+^ cells that originate from a non-SMC perivascular S100β^+^ parent population in support of other studies demonstrating a stem cell origin for lesional cells [46–48]. When the photonic profile of a range of cells *in vitro* and *ex vivo* were used as part of a training set by LDA, a substantial proportion of cells within lesions were classified as S100β^+^ vSCs and their SHh-derived myogenic progeny. The latter is not surprising as several previous studies support a role for Hh signalling in vascular lesion formation [19, 21, 22] where adventitial Sca1^+^ stem cells co-localise with SHh and its receptor, Ptch1 [49].

Multi-layered neural networks that mimic a human neural circuit structure have become increasingly popular in finding latent data structures and classifying highly nonlinear photonic datasets [13]. They have also proven ideal for classifying cell subtypes from label-free images [50] to predict macrophage activation [51], lymphocyte cell types [52], pluripotent stem cell-derived endothelial cells [53], and differentiating primary hematopoietic progenitors [54], Our analysis using a pre-trained MLP neural network clearly facilitated classification of cells from normal and injured vessels with autofluorescence emissions as the only input. Similarly, undifferentiated stem cells were easily discriminated from their myogenic progeny with the same trained photonic dataset. Using this MLP artificial neural network, we successfully classified sham and ligated cells *ex vivo* and further predicted the presence of S100β^+^ mVSc and their myogenic progeny within injured vessels confirming our S100β lineage tracing analysis and supporting previous studies that mapped stem cell-derived myogenic progeny to vascular lesions [46–48]. Our success in classifying these cells without exogenous contrast agents suggests that modern machine learning approaches may help compensate for spectral data with less molecular specificity and may provide further insight into the function and regulation of stem cells and their progeny in disease. Further enhancement of the AI model was facilitated when spectral signatures were combined with indices of cell metabolism, lineage specific gene expression and structural gene expression to predict the cellular heterogeneity within vascular lesions. Beyond the immediate findings of our study, these single-cell photonic profiles generated from healthy and injured vessels may also enable further examination of vascular cell heterogeneity and function in disease.

Mammalian cells are known to contain molecules which become fluorescent when excited by UV/Vis radiation of suitable wavelength arising from endogenous fluorophores [27], There are many endogenous fluorescence sources of contrast but the most robust and widely reported have been those associated with metabolism and structural architecture [55]. These signatures offer the greatest impact when deployed in combination with clinical assessment of normal, altered or diseased states and act as powerful intrinsic “optical antennas” of the morphological and functional properties of cells. [27], In stem cells, the autofluorescence signal is strongly dominated by cellular autofluorescence, for which NAD(P)H and FAD are the major contributors [59, 60], while other fluorophores such as collagens and elastin are dominant in vascular tissue [58]. The relative proportions of these fluorophores may change in disease and reflect the energetic and structural health of the vascular cells. While structural alterations are relatively straightforward to detect, due to the high quantum yield of collagen and elastin and their long fluorescence lifetime, metabolic alterations are more challenging to interpret due to the multitude of metabolic pathways and fluorophore species involved [31]. Vascular cells produce elastin as part of their reaction to increased mechanical stress associated with arteriosclerotic disease progression [59, 60] and it is significantly increased following carotid artery injury [28] and TGF-β1 stimulation [61]. Since specific knockdown of elastin caused a significant decrease in the photonic intensity of stem cells following myogenic differentiation, it is likely that changes in elastin content within ligated vessels due to the accumulation of stem cell-derived progeny may contribute to the enhanced emissions at this wavelength. In a similar manner, photonic emissions from collagens are generally associated with hydroxylysyl-pyridinoline and lysyl-pyridinoline groups that are affected by age-dependent polymerization of monomeric chains and cross links [62], In the normal arterial wall, collagen type I and Coll 3A1 are the main collagens in the media and adventitia while arterial injury alters the balance between TGF-β1 and these two types of collagen [63–65]. The deposition of Coll 3 has also been recently identified in neointimal SMC-like cells in a rat carotid model using 18F-fluorodeoxyglucose (FDG positron emission tomography – PET) [66]. Our study demonstrates that undifferentiated stem cells undergoing myogenic differentiation increase their collagen production concomitant with an increased photonic intensity at λ565nm. Since Col 3A1 emission spectra are observed at λ565 nm and Coll 3A1 depletion reduced this photonic intensity, it is likely that changes in collagen content within cells of ligated vessels due to the accumulation of stem cell-derived progeny may also contribute to the enhanced emissions at this wavelength.

Independent of structural changes, there is also recent evidence linking cell metabolism to transcriptional activity during differentiation/de-differentiation cycles that may contribute to changes in the photonic profile of these cells [67], Indeed, it is known that glucose catabolism due to cellular differentiation increases relative to oxidative phosphorylation during periods of biosynthesis with significant carbon demands [68]. In fully differentiated cells, such as medial SMCs, efficient ATP production is valuable, and accordingly, these cells primarily rely on oxidative phosphorylation, producing a high baseline optical redox ratio of FAD/[NAD(P)H+FAD] [71], While we clearly demonstrate upregulation of glycolytic metabolism in cells upon myogenic stimulation (even in the presence of oxygen) akin to the “Warburg effect”, the precise species involved were not specifically addressed in the current manuscript. NADPH is routinely utilized in the maintenance of pools of glutathione, thioredoxin, and peroxiredoxins, which help to create a reductive environment after oxidative damage produced by processes such as inflammation or ischemia, typical of vascular injury [72], Quantifying NAD(P)H and FAD fluorescence through an optical redox ratio and fluorescence lifetime imaging (FLIM) can provide sensitivity to the relative balance between oxidative phosphorylation and glucose catabolism. As peak single-photon NADH and NADPH fluorescence emissions are normally observed between λ440 and λ470⍰nm using λ330-360⍰nm excitation and FAD fluorescence is normally emitted at a peak of 520-530⍰nm when excited between 365 and 465⍰nm [73], it is unlikely that the changes at λ565 nm in cells from ligated vessels and S100β derived myogenic progeny are due to NAD(P)H emissions.

Importantly, programming and lineage-specific differentiation of vascular progenitors is associated with a metabolic phenotype [68–70]. Recent studies support such a concept since mitochondrial protein Poldip2 (Polymerase Delta Interacting Protein 2) controls SMC differentiation through a mechanism that involves regulation of metabolism [74], Poldip2 deficiency reduces the activity of the Krebs cycle and inhibits rates of oxidative metabolism while increasing rates of glycolytic activity in many different cell types, including vascular SMCs [67], While heterozygous Poldip2 deletion has no obvious phenotype, it is notable that mice are protected against neointimal formation following injury [75] underlying the importance of metabolism in disease progression. Therefore, it is possible that the changes in the photonic profile from ligated cells may also involve alterations in the metabolic state of these cells, in particular, if these cells are mVSc cell-derived myogenic progeny. While challenging to interpret, future studies will attempt to illuminate the potential contribution of individual key fluorophores involved in cell metabolism in single cells *ex vivo* during disease progression.

One important limitation of this study is the lack of an extensive library of spectral signatures or the use of nonlinear unmixing and other spectral analysis methods (which utilise similarity or spectral distance metrics) to estimate the abundances of autofluorescence molecules. Ideally, every source of fluorescence would be identified and its abundance quantified. While the impact of two types of fluorescent molecule known to be upregulated in vascular lesions [28, 66] was addressed, previous fluorescence imaging studies have shown significant spectral differences among the various types of collagens during fibrosis [9]. We clearly demonstrate that when recombinant Col 3A1 is excited at various wavelengths (λ300-500 nm), a clear emission spectra is present at λ465, λ530 and λ565 nm. It is therefore likely that the observed changes at λ565 nm in single cells from ligated vessels *ex vivo* and S100β derived myogenic progeny *in vitro* are due in part to increased Col 3A1 levels in these cells, in particular, since Col 3A1 depletion reduced the autofluorescence emissions at λ565 nm. Future work will address a wider range of autofluorescence molecules, including other types of collagen, lipofuscin and the contribution of NAD(P)H and FAD amongst others [77], Nevertheless, it is clear that endogenous fluorophores acting as intrinsic biomarkers offer an exceptionally powerful tool to characterize subtle changes of interconnected morphological and metabolic properties of cells and tissues under physiological or pathological conditions.

In conclusion, we have demonstrated the feasibility of single cell photonics to facilitate the detection of disease-relevant photonic signatures that reflect cell type and differentiation state. These signatures may have important predictive value for vascular disease and expand our current understanding of vascular pathology as they represent an important discriminator for classifying vascular phenotypes in subclinical atherosclerosis. As S100β^+^ mVSc and their myogenic progeny display a distinct photonic signature, this may facilitate early detection *in situ*. This approach is clinically feasible as the data are based on broadband light detection that can be implemented with state-of-the-art diagnostics using *in situ* endoscopic analysis [32] or integrated ultrasound and multispectral fluorescence lifetime catheters [76]. Our study supports implementation of a multifactorial approach in early vascular lesion detection where imaging and photonic signatures are combined to assess lesion progression thereby enabling more information about the disease before final treatment-decisions are made. Combining vascular phenotype information from photonic profiles with imaging may also be useful for other fibrotic diseases.

## Methods

### Biochip Device

The biochip design was based on cell sedimentation under stagnant flow conditions due to the application of centrifugal force into an array of V-shaped capturing elements (Figure 1E). The base (microfluidic inlets and V-cup array) section of the biochip was fabricated in PDMS (Sylgard 184, Dow Corning GmbH, Germany). Moulds for PDMS casting were surface micro-machined using SU8-3025 (Microchem, USA) for manufacturing the V-cup array and the reservoirs. The biochip middle layer (chip support holder) was manufactured using poly(methyl meth-acrylate) (PMMA) with a thin layer of pressure sensitive adhesive (PSA) attached to its base. A laser cutter (Epilog Zing Laser, Epilog, USA) defined the middle layer of the biochip. The chip substrate consisted of a standard borosilicate microscope slide which was bonded to the chip middle layer using PSA. This hybrid chip was then treated by air plasma (1000 mTorr) for 5 minutes and assembled with the PDMS base to complete the biochip [78] The fabrication method for the biochip ensured transparency and biocompatibility and was leak free and facilitated long term stability. The operating principles of the V-cup array have been described previously [11]. Briefly, the sedimentation takes place with the liquid bulk at rest to provide high capture efficiency. V-cups (13 μm diameter) staggered in an array of 47 x 24 cups which can thus trap up to 1128 individual cells. Additional trap and pillar based locations are also on the biochip to facilitate a sub-population of cells to be selected (via optical tweezers) and further single cell assays. Finally, a disc for holding three biochips and for mounting onto the centrifugal test stand was manufactured using 3D printing. The centrifugal test setup comprised a motor for spinning the microfluidic chips (4490H024B, Faulhaber micromotor SA, Switzerland), a synchronized camera for image acquisition during rotation (TXG14c, Baumer AG, Germany) coupled to a motorized 12x zoom lens (Navitar, USA) and a strobe light unit (Drelloscop 3244, Drello, Germany) as described previously [78]. The system integration between the microfluidic and optical systems for SCA was performed using an in-built optical detection and imaging system on the centrifugal test stand. The optical module incorporated a laser tweezers to manipulate individual cells on disc using a 1-W, 1064-nm infrared laser (Roithner Lasertechnik, Austria). This laser was focused through a 40x oil immersion microscope objective (CZ Plan Neofluar 40x/1.3 OIL PH3, Zeiss, Germany) with a numerical aperture (NA) of 1.3. This setup allowed a working distance of 200 μm. This objective was mounted on a piezo driven Z-drive with a travel range of 100 μm (Fast PIFOC^®^ Piezo Nanofocusing Z-Drive, PI, Germany) for fine focusing. Additionally, the module included a high sensitivity cooled CCD camera (Sensicam qe, PCO, Germany) which utilizes the same optical path as the laser to facilitate particle handling and ac-quisition of bright-field and fluorescent images. Excitation was performed by a 250-W halo-gen lamp (KL 2500 LCD, Schott, Germany) with an enclosed filter wheel to allow both broad-band light (λ = 360 – 800 nm) and selected fluorescent excitation (filtered at excitation wave-lengths of 403 ± 32 nm, 492 ± 15 nm, 515 ± 25 nm, 572 ± 15 nm and 610 ± 32 nm) and emission wavelengths (emission filters are 465 ± 20 nm, 530 ± 20 nm, λ565 ± 20 nm and 630 ± 20 nm and 670 ± 20 nm). The module was mounted on a computer controlled X-Y stage (Qioptiq, Germany). Measurement of background photons assessed the contribution of the chip PDMS material and the surrounding liquid (cell media), on a fully primed empty chip. The images are acquired post capture when the cells settle into the capture V-cup region. The method of travel through the chip was sedimentation based and did not impact on the fluorescence emissions as the cells experience minimum shear force as they travel through the chip. Spin speed for the cells travelling through the chip is fixed at 10Hz for all experiments to reduce any instrumentation variations. Excitation (lamp power etc.) and image acquisition settings (exposure time, min and max values etc.) were fixed and uniform across all experiments. The effect of the width of the excitation (15-32 nm) using commercial filters fitted into the imaging system revealed that changing an excitation filter bandwidth by a small increments had little impact on fluorescence excitation. In all studies, the entire cell was selected as the region of interest (ROI) and the mean value across the entire cell is presented. The experimental volume fraction of the cell probed and the standard deviation across each cell, in each wavelength, was low (data not shown).

### Biochip Preparation and Microfluidic Testing

The device was placed in vacuum prior to introducing the liquids for a minimum of 30 minutes ensure complete and bubble-free filling. The biochip was primed with the appropriate cell culture media formulation for the cell type under test via the loading chamber on the top right section of the biochip. After priming, cells were introduced via the loading chamber on the top left section of the biochip. All pumping was performed the centrifugal test stand and a 3D-printed chip holder which allows three biochips to be tested in parallel, thus significantly increasing the cell capture efficiency of the V-cup array system compared to common, flow-driven systems [78]. Initial cell capture tests were performed using 20-μm polystyrene beads to emulate cell behaviour before being repeated using cells. In all cases, the sedimentation in absence of flow led to significantly increased occupancy of the V-cups (≥ 95%) compared to common, flow-driven methods (data not shown).

### Tamoxifen-induced genetic lineage tracing

*S100β-CreER*-Rosa26tdT mice (average weight 20g, 6-8 wks old) were injected IP with tamoxifen (Tm) dissolved in corn oil at 75 mg/kg for 5 consecutive days. Carotid artery ligation (partial, or complete), or sham operation, was performed at 4 weeks after the last injection of Tm. At the indicated time post-ligation or sham operation, anesthetized mice were perfusion fixed and carotid arteries harvested for analysis.

### Histomorphometry

At specified times post-ligation, mice were anesthetized (ketamine/xylazine) and perfusion fixed with 4% paraformaldehyde in sodium phosphate buffer (pH 7.0). Fixed carotids were embedded in paraffin for sectioning. Starting at the carotid bifurcation landmark (single lumen) a series of cross-sections (10 x 5 μm) were made, every 200 μm through 2 mm length of carotid artery. Cross-sections were de-paraffinized, rehydrated in graded alcohols and stained with Verhoeff-Van Gieson stain for elastic laminae and imaged using a Nikon TE300 microscope equipped with a Spot RT digital camera (Diagnostic Instruments). Digitized images were analyzed using SPOT Advanced imaging software. Assuming a circular structure *in vivo*, the circumference of the lumen was used to calculate the lumen area, the intimal area was defined by the luminal surface and internal elastic lamina (IEL), the medial area was defined by the IEL and external elastic lamina (EEL), and the adventitial area was the area between the EEL and the outer edge, essentially as described by us previously [22],

### Haematoxylin and Eosin Staining of Tissue

For murine vessels, paraffin rehydration was conducted at room temperature by immersing the slides in xylene for 20 minutes. The slides were then immersed in the following gradients of ethanol for 5 minutes respectively: 100 %, 90 %, 70 % and 50 % ethanol. The slides were then rinsed in distilled H_2_O_2_ before washing in 1x PBS for 10 minutes. The slides were stored in 1x PBS until ready when they were immersed in Harris Haematoxylin at room temperature for 8 min.

### Immunofluorescent Staining of tissues

Immunostaining essentially as previously described [22], Carotid artery cryosections were air-dried for 1 h at room temperature, followed by incubation with blocking buffer (5% donkey serum, 0.1% Triton X-100 in PBS) for 30 minutes at room temperature and then incubated with primary antibody overnight at 4°C in antibody solution (2.5% BSA, 0.3M Glycine and 1% Tween in DPBS). Murine and human arterial sections were stained with primary antibodies [Supplementary Table I]. Isotype IgG control and secondary antibody only controls were performed. For antigen retrieval, slides were brought to a boil in 10 mM sodium citrate (pH 6.0) then maintained at a sub-boiling temperature for 10 minutes. Slides were cooled on the bench-top for 30 minutes then washed in deionized water (3 × 5 min) each before being washed in PBS (3×5 min). The antigen retrieval protocol diminishes endogenous eGFP and Tdt tomato transgene signals. Therefore, those sections were costained with anti-eGFP antibody and anti-Td tomato antibody [Supplementary Table I].

For immunofluorescence staining, 5 consecutive images were obtained and processed using ImageJ software^™^ to analyze the collected images. Images were merged using the Image color-merge channels function. Merged signals and split channels were used to delineate the signals at single-cell resolution. Settings were fixed at the beginning of both acquisition and analysis steps and were unchanged. Brightness and contrast were lightly adjusted after merging.

### Immunocytofluorescent staining of cells

Cells seeded onto UV sterilized coverslips were fixed with 3.7 % formaldehyde, (15 min, RT). If cells required permeabilization for the detection of intracellular antigens, cells were incubated in 0.025 % Triton X-100 PBS (room temp, 15 min). All coverslips were blocked (1 hr, RT) using 5 % BSA, 0.3 M Glycine, 1 % Tween PBS solution (1 hr, RT). Cells were incubated overnight with primary antibodies at 4°C [Supplemental Table II], then washed twice with PBS to remove any unbound primary antibody before being incubated (1 hr, RT) with the recommended concentration of fluorochrome-conjugated secondary antibodies diluted in blocking buffer [Supplementary Table II]. Following 2x wash in PBS, cell nuclei were stained using DAPI: PBS (dilution 1:1000) (15 min, RT). For each primary and secondary antibody used, a secondary control and an IgG isotype control was performed to assess nonspecific binding. An Olympus CK30 microscope and FCell^™^ software was used to capture images. Images were analysed using ImageJ software as described above. Settings were fixed at the beginning of both acquisition and analysis steps and were unchanged. Brightness and contrast were lightly adjusted after merging.

### Chromatin Immunoprecipitation (ChIP)

ChIP was performed on cultured cells as previously described with slight modifications [79] Protein-DNA interactions were cross-linked in cells at 70% confluency with 1% paraformaldehyde (10 min, RT). Cross-linking was stopped by addition of 125 mM glycine for 10 min and the chromatin was sonicated to shear chromatin into fragments of 200-600 base pairs. The sheared chromatin was immunoprecipitated with 2 μg tri-methyl-histone H3 (Lys27) and di-methyl-histone H3 (Lys4), while negative control was incubated with mouse IgG and input DNA without antibody using the CHIP—IT Express HT Kit from Active Motif (Cat no: 53018) according to the manufacturer’s instructions [Supplementary Table III]. A chromatin IP DNA Purification Kit (Cat No 58002 Active Motif) was used to purify the samples after CHIP before PCR was performed using Myh11 promoter primers [5’ -CCC TCC CTT TGC TAA ACA CA – 3′ and 5’ – CCA GAT CCT GGG TCC TTA CA – 3] as previously published [79]. Sensimix SYBR^®^ no-ROX Bioline Kit (QT650) was used to perform Real Time One-Step PCR according to the manufacturers’ instructions. The antibodies used for ChIP are outlined in Supplemental Table I.

### Quantitative PCR

Total RNA was prepared from cultured cells using the ReliaPrep^™^ RNA Cell Miniprep System kit from Promega according to the manufacturer’s protocol. Two micrograms of RNA was used for reverse transcription with Rotor-Gene SYBR Green RT-PCR (QIAGEN) or The SensiMix^™^ SYBR^®^ No-ROX (BioLine) protocols for Real TimeOne-Step RT-PCR using the Real Time Rotor-GeneRG-3000^™^ light cycler from Corbett Research using primers listed in Supplementary Table IV.

### Western blot analysis

Total protein (~40μg) was resolved with SDS-PAGE and transferred to nitrocellulose membranes. The membrane was probed with primary antibodies [Supplementary Table II] and secondary anti-rabbit-anti-mouse antibody, HRP conjugated A5278 (Sigma) and antirabbit antibody, HRP conjugated A0545 (Sigma). Detection was performed using TMB T0565 (Sigma).

### Collagen autofluorescence

The level of collagen autofluorescence emissions was measured using recombinant 100ng/ml Col 1⍰1, Col 1⍰2 and Col 3A1 (Cat Nos abx065998, abx167199 and abx066003, Abbexa, Cambridge UK) suspended in PBS using the Load platform. Parallel experiments were conducted using a Tecan Infinite 200 model multifunctional plate reader. Background levels of emissions due to PBS were subtracted before spectra were analysed using Prism GraphPad Software, v9.

### Cell Metabolism Assays

The level of glucose, glutamine and lactate was measured in conditioned media from undifferentiated stem cells before and after myogenic differentiation with Jag-1 (1.0 μg/ml). L-Lactate and Glutamine was measured using commercial assay kits K-GLNAM-L-Glutamine/Ammonia Assay Kit (Rapid) from Megazyme (Megazyme Ltd, Ireland) and Lactate Assay II kit from Sigma (Merck KGaA, Darmstadt, Germany), according to the manufacturer’s instructions. Glucose consumption was measured using GlucCell Glucose Test Strips on a GlucCell glucose monitoring system (KDBio, UK).

## QUANTIFICATION AND STATISTICAL ANALYSIS

All data were determined from multiple individual biological samples and presented as mean values ± standard error of the mean (SEM). All *in vitro* experiments were performed in triplicate and repeated three times unless otherwise stated. All mice were randomly assigned to groups (including both male and female), and the analysis was performed blind by two groups. For statistical comparisons, all data was checked for normal gaussian distribution before parametric and non-parametric tests were performed. An unpaired twosided Student’s t-test was performed to compare differences between two groups and significance was accepted when p≤ 0.05. A one sample t Wilcoxon test was performed on fold changes and significance was accepted when p≤ 0.05. An ANOVA test was performed for multiple comparisons with a Sidak’s multiple comparisons test for parametric data and a Dunn’s Multiple comparison test for non-parametric data using GraphPad Prism software v8^™^. and significance was accepted when p≤ 0.001.

For PCA and LDA multivariate analysis, the cells were first classified to create a ground truth for each sample before the spectral data were normalized to one by dividing the fluorescence intensity at each wavelength by the average background intensity at that wavelength. Dimensionality reduction was achieved through a Multiclass Fisher’s Linear Discriminant Analysis (Multiclass FLDA) preprocessing filter, as previously described using WEKA machine learning tool kit, version 3.8.4 [80] and further analyzed by PCA and LDA using the multivariate statistical package, PAST4. A scatter plot of specimens along the first two canonical axes produces maximal and second to maximal separation between all groups. The axes are linear combinations of the original variables as in PCA, and eigenvalues indicate amount of variation explained by these axes. When only two groups are analyzed, a histogram is plotted. The data are classified by assigning each point to the group that gives minimal Mahalanobis distance to the group mean. The Mahalanobis distance is calculated from the pooled within-group covariance matrix, giving a linear discriminant classifier. The given and estimated group assignments are listed for each point. In addition, group assignment is cross-validated by a leave-one-out cross-validation (jackknifing) procedure. To interrogate the photonic profiles from unknown ‘mystery’ groups, i.e. data that are not included in the discriminant analysis itself but are classified. In this way, it is possible to classify new datasets that are not part of the training set.

### Supervised machine learning (ML)

Supervised ML is a technique in which a model is trained on a range of inputs (or features) which are associated with a known outcome. Once the algorithm is successfully trained, it will be capable of making outcome predictions when applied to new data. Spectra from single cells across five broadband light wavelengths were first classified to create a ground truth for each sample before the data were normalized as before. Dimensionality reduction was achieved through Multiclass Fisher’s Linear Discriminant Analysis (Multiclass FLDA), as previously described before MLP artificial neural network analysis was performed on each dataset using the WEKA machine learning tool kit [81]. Once the dataset was loaded and pre-processed using the FLDA filter, it was processed using the “Classify” panel with the implementation for MLP artificial neural network (ANN) analysis. ANNs are algorithms which are loosely modelled on the neuronal structure observed in the mammalian cortex and are arranged with a number of input neurons, which represent the information taken from each of the features in the dataset which are then feed into any number of hidden layers before passing to an output layer in which the final decision is presented. The algorithm was iteratively improved using an optimization technique by changing the number of hidden layers (1,2.3., etc) and the momentum (0.1-0.5) and rate of learning (0.1-0.5) to reduce the error of prediction and then evaluated using cross-validation. The risk of over-fitting was mitigated against when our dataset was split into two segments; a training segment and a testing segment to ensure that the training model can generalize to predictions beyond the training sample. Each segment contained a randomly selected proportion of the features and their related outcomes which allowed the algorithm to associate certain features, or characteristics, with a specific outcome, and is known as training the algorithm. Once training was completed, the algorithm was then applied to the features in the testing dataset without their associated outcomes. The predictions made by the algorithm were then compared to the known outcomes of the testing dataset to establish model performance. A total of 924 cells were used to create this dataset across the five wavelengths. The initial model algorithm had 2 hidden layers and a momentum rate of 0.2 and a learning rate of 0.3. When the additional variables of SMC gene expression, glycolytic metabolism and Col 3A1 were included, the model algorithm had 1 hidden layer and a momentum rate of 0.2 and a learning rate of 0.3. Performance was evaluated based on F1 scores, an aggregate of recall and precision, and on accuracy, the fraction of correct predictions.

## EXPERIMENTAL MODEL AND SUBJECT DETAILS

### Mice Breeding and Genotyping

All procedures were approved by the University of Rochester Animal Care Committee in accordance with the guidelines of the National Institutes of Health for the Care and Use of Laboratory Animals. S100β -EGFP/Cre/ERT2 transgenic mice (JAX Labs, stock #014160, strain name B6;DBA-Tg(S100β -EGFP/cre/ERT2)22Amc/j) express the eGFPCreER^T2^ (Enhanced Green Fluorescent Protein and tamoxifen inducible cre recombinase/ESR1) fusion gene under the direction of the mouse S100β promoter. Ai9 mice (Jax Labs, stock #007909, strain name B6.Cg-Gt(ROSA)26Sor^tm9(CAG-tdTomato)Hze^ /J) express robust tdTomato fluorescence following Cre-mediated LoxP recombination. For lineage tracing experiments S100β-eGFP/Cre/ERT2-dTomato double transgenic mice of both genders were generated by crossing S100β-eGFP/Cre-ERT2 mice with Ai9 reporter mice. The tdTomato transgene expression pattern corresponds to genomically marked S100β and the eGFP transgene expression pattern corresponds to current expression of S100β. Genomic DNA for genotyping was prepared from mice tail and papered for genotyping by Proteinase K lysed, isopropanol precipitated and 70% ethanol washed. The number of animals used were approved based on the experiments effects size. All male and female mice were included in the study and were 8-10 weeks old.

### Carotid Artery Ligation

Ligation of the left common carotid artery was performed four weeks after Tm-induced cre-recombination in randomised male and female S100β-eGFP/Cre/ERT2-dTomato double transgenic mice, essentially as described previously [22], Mice that remained healthy and had a pronounced intimal lesion after 21 days were included. Prior to surgery mice received a single dose of Buprenorphine SR (sustained release) analgesia (0.5-1.0 mg/kg SQ) (and every 72 hrs thereafter as needed). The animal was clipped and the surgical site prepped using betadine solution and alcohol. A midline cervical incision was made. For partial carotid artery ligation, with the aid of a dissecting microscope, the left external and internal carotid arterial branches were isolated and ligated with 6-0 silk suture reducing left carotid blood flow to flow via the patent occipital artery. The neck incision (2 layers, muscle and skin) was sutured closed. Partial ligation of the left carotid artery in this manner resulted in a decrease (~80%) in blood flow, leaving an intact endothelial monolayer. Buprenorphine was administered at least once post-op 6-12 hrs. For complete ligation, the left common carotid was isolated and ligated just beneath the bifurcation with 6-0 silk suture. In sham surgery group, carotid was similarly manipulated but not ligated. The neck incision (2 layers, muscle and skin) was sutured closed and the animal allowed recover under observation. After 14 and 21 days, mice were perfused with 4% PFA via the left ventricle for 10 min and carotid arteries were harvested for morphometric and histochemical analysis.

In parallel studies, cells from mouse carotids and thoracic aorta were isolated and placed in cold Hank’s solution for adipose tissue removal, lumen rinsing and removal of the endothelium by gentle rubbing. The adventitia was enzymatically removed by incubation of the vessels in collagenase solution [i.e., MEM⍰ nucleosides GlutaMAX (2 mL) containing Collagenase type 1A (0.7 mg/mL), soybean trypsin inhibitor (50 mg/mL), and bovine serum albumin (1 mg/mL)] for approx. ~ 20 min at 37°C. Once the adventitia became lose, it was carefully removed as an intact layer using forceps under a dissecting microscope. The aorta was cut into 1 mm pieces and digested with Elastase type III (Sigma) at 37°C before the dispersed cells were centrifuged, washed twice in Hanks Balanced salt solution and fixed with 3.7% PFA before spectral analysis on the LoaD platform.

### Cell culture and harvesting

Murine aortic SMCs (Movas (ATCC^®^ CRL-2797^™^) were cultured in Dulbeco’s Modified Eagles growth medium (DMEM) supplemented with 10% Fetal Bovine Serum, 2 mM L-glutamine) and 1% penicillin-streptomycin. Murine embryonic C3H/10T1/2 cells (ATCC^®^ CRL-226^™^) were grown in Eagle’s Basal medium supplemented with heat-inactivated fetal bovine serum to a final concentration of 10% and 2mM L-glutamine and 1% penicillin-streptomycin. Murine vSCs from mouse aorta were grown in maintenance media (MM) containing DMEM media supplemented with chick embryo extract (2%), FBS embryonic qualified (1%, ATCC), B-27 Supplement, N-2 Supplement (Cell Therapy Systems), recombinant mouse bFGF (0.02 μg/mL), 2-Mercaptoethanol (50 nM), retinoic acid (100 nM), Penicillin-Streptomycin (1%). J774A.1 mouse BALB/c monocyte macrophages were a kind gift from Prof Christine Loscher, DCU and were grown in DMEM supplemented with 10% Fetal Bovine Serum, 2 mM L-glutamine and 1% Penicillin-Streptomycin. Human Burkitt’s lymphoma B cells (Ramos B cells) were a kind gift from Prof Dermot Walls, DCU and were grown in RPMI 1640 supplemented with 10% Fetal Bovine Serum (Heat inactivated), and 1% penicillinstreptomycin.

### Isolation of S100β/Sca1_+_ murine resident vascular stem cells (vSCs)

Using an optimised cell dissociation protocol, mVSc from mouse aorta were isolated using sequential seeding on non-adherent and then adherent plates. Mouse thoracic aortas (4 at a time) were harvested and placed in cold Hank’s solution for adipose tissue removal and lumen rinsing. The adventitia was enzymatically removed by incubation of the vessels in collagenase solution [i.e., MEM⍰ nucleosides GlutaMAX^™^ (2 mL) containing Collagenase type 1A (0.7 mg/mL), soybean trypsin inhibitor (50 mg/mL), and bovine serum albumin (1 mg/mL)] for approx. 10-20 min at 37°C. Once the adventitia became lose, it was carefully removed as an intact layer using forceps under a dissecting microscope. The aorta was cut into 1 mm pieces and digested with Elastase type III (Sigma) at 37°C. Dispersed cells were centrifuged and washed twice in warm maintenance medium (MM) (DMEM supplemented with 2% chick embryo extract, 1% FBS, 0.02 μg/mL, bFGF basic protein, B-27 Supplement, N-2 supplement, 1% Penicillin-Streptomycin, 50 nM 2-Mercaptoethanol and 100 nM retinoic acid) before seeding (1st seeding) on a 6-well non-adherent plate in MM. At 48 h, suspension cells were transferred (2nd seeding) to a fresh well of a CELLstart^™^ pre-coated 6-well plate in MM. MM was added to the remaining cells in the non-adherent surface. Cells were incubated at 37°C, 5% CO_2_ for 1 week with minimal disturbance. The stem cells exhibited a uniform neural-like morphology in low density culture adopting a dendritic-like tree shape and retaining their morphological characteristics at low density throughout repeated passage. Cells were fed with MM every 2-3 days and passaged every 3-4 days or when ~70% confluent.

## Declarations

### Funding (information that explains whether and by whom the research was supported)

This research was funded in part by Science Foundation Ireland grant SFI-11/PI/1128, Health Research Board (HRB) of Ireland grant HRA-POR-2015-1315, and the European Union’s INTERREG VA Programme, managed by the Special EU Programmes Body (SEUPB) to PAC, and NIH R21AA023213 and RO1AA024082 to EMR, Irish Research Council (IRC) GOIPG/2014/43 (M.DiL), Fraunhofer-Gesellschaft under the SFI Strategic Partnership Programme [Grant Number 16/SPP/3321] Science Foundation Ireland [Grant Number: 10/CE/B1821]; the ERDF; the LiPhos project funded by the European Commission [Grant Number: 317916] to JD.

### Conflicts of interest/Competing interests (include appropriate disclosures)

The authors declare no competing financial interests or conflicts of interest.

### Ethics approval (include appropriate approvals or waivers)

Part of the animal studies were approved by The Jackson Laboratory Animal Care and Use Committee (Permit Number: 07007) and were in accordance with the “Guide for the Care and Use of Experimental Animals” established by the National Institutes of Health (1996, revised 2011). Part of the animal studies were also approved by the University of Rochester Animal Care Committee in accordance with the guidelines of the National Institutes of Health for the Care and Use of Laboratory Animals.

### Consent for publication (include appropriate statements)

All authors have given their consent for publication and have reviewed and approved the submission.

### Authors’ contributions

E.M.R, D.M., and W.L., performed the animal experiments and M.DiL performed the blind analysis of the confocal images. C.M., M.DiL., R.H., D.B., E.F., performed the murine cell culture experiments, D.B., MDiL, Y.G., and R.H., performed the ChIP analysis. D.K., A.L., L.J., and J.D. developed the LoaD platform and C.M, A.O., D.K, D.C., K.H., and R.H performed the single cell photonic analysis studies. M.K performed the collagen spectroscopy and metabolic assays. E.F and P.A.C performed the PCA/LDA and WEKA analysis. E.M.R., C.M., and P.A.C drafted the manuscript, revised the manuscript and discussed the results. E.M.R., D.K., CM and P.A.C., designed and coordinated the experiments, interpreted the data, revised and confirmed the paper. All authors reviewed and confirmed the manuscript.

### Availability of data and material (data transparency)

Several datasets were generated and analysed during the current study and will be made available on reasonable request.

## Acknowledgments

We thank Diana Scott for histological tissue processing, and Drs Linda Callahan and Paivi Jordan (University of Rochester, NY) for confocal microscope expertise and Dr Tim Downing (Dublin City University) for the advice on the computational and statistical analysis. We thank Prof Robert Forster and Dr Phil Cummins for critically reviewing the manuscript and for their insights and helpful discussions.

