## Supplementary Material for "Disease-relevant single cell photonic signatures identify S100β stem cells and their myogenic progeny in vascular lesions"

Supplementary Table I

Antibodies and their corresponding dilutions used in immunohistochemistry.

| **Antibody/Product Name** | **Supplier/Product Number** | **Dilutions** |
| --- | --- | --- |
| Rabbit anti-mouse/rat alpha smooth muscle cell ⍺-actin antibody | Abcam (ab5694) | 1/200 |
| Anti-Actin, α-Smooth Muscle antibody, Mouse monoclonal | Sigma (A5228) | 1/200 |
| Anti-S100-β (CT) Antibody, clone EP1576Y, rabbit monoclonal | Millipore (04-1054) | 1/100 |
| Chicken anti-GFP antibody | Abcam (ab13970) | 1/1000 |
| Rabbit Anti-RFP/dT antibody | Abcam (ab62341) | 1/1000 |

Supplementary Table II

Antibodies and their corresponding dilutions used in immunocytochemistry/western blot.

| **Antibody/Product Name** | **Supplier/Product Number** | **Dilutions** |
| --- | --- | --- |
| Rabbit anti-mouse Calponin [EP798Y] | Abcam (ab46794) | 1/200 |
| Goat anti-mouse/rat/human smooth muscle Myosin heavy chain | Santa Cruz (sc-79079) | 1/200 |
| Rabbit Anti-S100β | Merck Millipore (ABN59) | 1/100 |
| Rabbit Anti-mouse/rat S100β [EP1576Y] | Abcam (ab52642) | 1/100 |
| Alexa Fluor® 488 Goat anti-mouse IgG | Invitrogen (A-11001) | 1/1000 |
| Alexa Fluor® 488 Goat anti-rabbit IgG | Invitrogen (A-11008) | 1/1000 |
| Alexa Fluor® 488 Donkey anti-goat IgG | Invitrogen (A-11055) | 1/1000 |

Supplementary Table III

**Antibodies used in Chromatin Immunoprecipitation (ChIP)**

| **Antibody/Product Name** | **Supplier/Product Number** |
| --- | --- |
| Rabbit anti-mouse Tri-Methyl-Histone H3 (Lys27) [C36B11] | Cell Signalling Technology (9733S) |
| Rabbit anti-mouse Di-Methyl-Histone H3 (Lys4) [C64G9] | Cell Signalling Technology (9725S) |
| Normal Rabbit IgG (ChIP graded) | Cell Signalling Technology (2729) |

Supplementary Table IV

Customised primers used in this study from Integrity DNA Technology (IDT).

| **Customised primer** | **Sequences** | |
| --- | --- | --- |
| Mm_Sm-mhc (Myh11) | Forward | 5' - GCA GTG AGC TCT CAG TCA TC - 3’ |
|  | Reverse | 5' - CAA TGC CTC CTC TGA CAA GT - 3' |
| Mm_Cnn1 | Forward | 5’ - GCT TGT CTG CTG AAG TAA AGA AC - 3' |
|  | Reverse | 5’ - TCC ATG AAG TTG TTC CCG ATG - 3' |
| Mm_Hprt | Forward | 5’ - GGC TAT AAG TTC TTT GCT GAC CTG C - 3' |
|  | Reverse | 5’ - GCT TGC AAC CTT AAC CAT TTT GGG - 3' |
| Mm_Gapdh | Forward | 5’ - GCC TCC AAG GAG TAA GAA AC - 3' |
|  | Reverse | 5’ - GCC TCC AAG GAG TAA GAA AC - 3' |
| Mm_ Sm-mhc (Myh11) for ChIP PCR | Forward | 5’ - CCC TCC CTT TGC TAA ACA CA - 3' |
|  | Reverse | 5’ - CCA GAT CCT GGG TCC TTA CA - 3' |

Primers used in this study from QIAGEN.

| **Primer** | **Product Name** | **Product Code** |
| --- | --- | --- |
| mHprt-1 | Mm_Hprt_1_SG QuantiTect Primer Assay | QT00166768 |
| mGapdh | Mm_Gapdh_3_SG QuantiTect Primer Assay | QT01658692 |
| mCol3A1 | Mm_Col3A1_3_SG QuantiTect Primer Assay | QT01055516 |

**Supplemental Figure Legends**

Figure S1. **Single cell photonics of Ramos B and J774A.1 macrophages. A.** Visualisation of Ramos B and J774A.1 cells on each V-cup in the LoaD platform. **B.** PCA loading plots of Ramos B and J774A.1 cells. **C.** LDA of Ramos B and J774A.1 cells. Data are from 55 cells/group over five wavelengths. **D-G.** Single cell auto-fluorescence photon emissions across five wavelengths from (D, F) Ramos B cells and (E, G) J774A.1 cells *in vitro* compared to sham (D, E) and ligated cells (F, G) *ex vivo*. Data are the Log2 fold increase and represent the mean ± SEM of 55-178 cells/group, #p≤0.001 vs Sham (D,E) Ligated (F,G). **H**. PCA loading plots of sham (black), ligated (orange), Ramos B (red) and J774A.1 (blue) cells. **I**. LDA plots of sham (black), ligated (orange) cells ex vivo and Ramos B (red) and J774A.1 (blue) cells in vitro. Data are from 466 cells across the five wavelengths **J**. Confusion matrix of true class and predicted class following a leave-one-out cross-validation procedure by the LDA classifier.

Figure S2. **Single cell photonics of MSC cells and their myogenic progeny compared to aortic SMC, Movas SMC, Ramos B cells and J774A.1 macrophages *in vitro*. A.** PCA loading plots of MSCs and their myogenic progeny, Ramos B cells and J774A.1 macrophages. Data are from 55-79 cells/group. **B.** LDA plot of MSCs and their myogenic progeny, Ramos B cells and J774A.1 macrophages. **C.** PCA loading plots of MSCs and their myogenic progeny, aortic SMCs and Movas SMCs. **D.** LDA plots of MSCs and their myogenic progeny, aortic SMCs and Movas SMCs. Data are from 55-79 cells/group. **E.** Confusion matrix of true class and predicted class following a leave-one-out cross-validation procedure using the LDA classifier. Data are from 378 cells across five wavelengths. **F.** Confusion matrix of true class and predicted class following a leave-one-out cross-validation procedure using the LDA classifier. Data are from 378 cells across five wavelengths.

Figure S3. **Single cell photonics of C3H10T1/2 cells and their myogenic progeny compared to Ramos B, J774A.1, aortic SMC, Movas SMC and MSC and their myogenic progeny *in vitro*. A.** PCA loading plots of C3H10T1/2 cells and their myogenic progeny compared to Ramos B cells and J774A1 cells. **B** LDA plots of C3H10T1/2 cells and their myogenic progeny compared to Ramos B cells and J774A1 **C.** PCA loading plots of C3H10T1/2 cells and their myogenic progeny, aortic SMCs and Movas SMCs. Data are from 55 cells/group. **D.** LDA plots of C3H10T1/2 cells and their myogenic progeny, aortic SMCs and Movas SMCs. **E.** PCA loading plots of C3H10T1/2 cells and their myogenic progeny, MSCs and their myogenic progeny. Data are from 55-79 cells/group. **F.** LDA plots of C3H10T1/2 cells and their myogenic progeny, MSCs and their myogenic progeny. **G.** Confusion matrix of true class and predicted class following a leave-one-out cross-validation procedure using the LDA classifier. Data are from 220 cells across five wavelengths. **H.** Confusion matrix of true class and predicted class following a leave-one-out cross-validation procedure using the LDA classifier. Data are from 367 cells across five wavelengths.

Figure S4. **Single cell photonics of mVSc and their myogenic progeny compared to aortic SMC, Movas SMC, Ramos B and J774A.1 macrophages. A.** Representative immunocytochemical analysis of S100β expression in mVSc. **B.** Visualisation of mVSc and their myogenic progeny on each V-cup in the LoaD platform. **C.** PCA loading plots of mVSc and their myogenic progeny, aortic SMCs and Movas SMCs. Data are from 55 cells/group. **D.** LDA plot of mVSc and their myogenic progeny, aortic SMCs and Movas SMCs. Data are from 55 cells/group. **E.** Confusion matrix of true class and predicted class following a leave-one-out cross-validation procedure using the LDA classifier. Data are from 319 cells across five wavelengths. **F.** PCA loading plots of mVSc and their myogenic progeny, Ramos B cells and J774A.1 macrophages. Data are from 55 cells/group. **G.** LDA plot of mVSc and their myogenic progeny, Ramos B cells and J774A.1 macrophages. Data are from 55 cells/group. **H.** Confusion matrix of true class and predicted class following a leave-one-out cross-validation procedure using the LDA classifier. Data are from 319 cells across five wavelengths.

Figure S5. **Single cell photonics of mVSc and their myogenic progeny compared to MSCs and their myogenic progeny. A.** PCA loading plots of mVSc and their myogenic progeny, and MSC and their myogenic progeny. Data are from 55-79 cells/group. **B.** LDA plot of mVSc and their myogenic progeny, and MSC and their myogenic progeny. Data are from 55-79 cells/group. **C.** Confusion matrix of true class and predicted class following a leave-one-out cross-validation procedure using the LDA classifier. Data are from 487 cells across five wavelengths.

Figure S6. **Single cell diameter measurements of cells isolated from sham and ligated vessels *ex vivo,* and aortic SMCs, Ramos B cells and myogenic progeny *in vitro*. A.** Diameter measurements of cells isolated from sham and ligated vessels captured on V-cups compared to aortic SMCs. **B.** Diameter measurements of MSCs before and after myogenic differentiation with TGF-β1 for 14d. **C.** Diameter measurements of C3H 10T1/2 cells before and after myogenic differentiation with TGF-β1 for 14 d. **D.** Diameter measurements of Ramos B cells before and after treatment with TGF-β1 for 14 d. Data are from 25 cells/group.

Figure S7. **The effect of myogenic differentiation on S100β stem cell metabolism. A.** The level of glucose, **B.** glutamine and **C.** lactate release in conditioned media from S100β mVSc before and after treatment with the myogenic stimulus, Jag-1 (1μg/ml) over time. **D-F.** The level of glucose consumption, glutamine consumption and lactate release in conditioned media after treatment with Jag-1 (1μ/ml) in the absence or presence of Notch inhibitor , DAPT (100μM). *p<0.05 vs Fc Control, # p<0.05 vs Jag1. Data are mean ± SEM of three individual wells.

Figure S8. **Collagen autofluorescence. A**. Representative autofluorescence photon emissions from recombinant Col 1⍺1, Col 1⍺2, and Col3A1 across five wavelengths using the Load platform. Date are corrected for background emissions from media. Data are the Log2 fold increase and represent the mean of three samples **B-F.** Autofluorescence emission spectra from Col 1⍺1, Col 1⍺2, and Col3A1 following excitation at λ358- 565 nm. Data are representative of two independent experiments. **G-I.** Autofluorescence emission spectra from Col3A1 at λ465, λ530 and λ565 following excitation at λ300- 500 nm. Data are representative of two independent experiments. Date are corrected for background emissions from PBS.
