## Supplementary figures and images for "Disease-relevant single cell photonic signatures identify S100β stem cells and their myogenic progeny in vascular lesions"

### Supplementary Figures

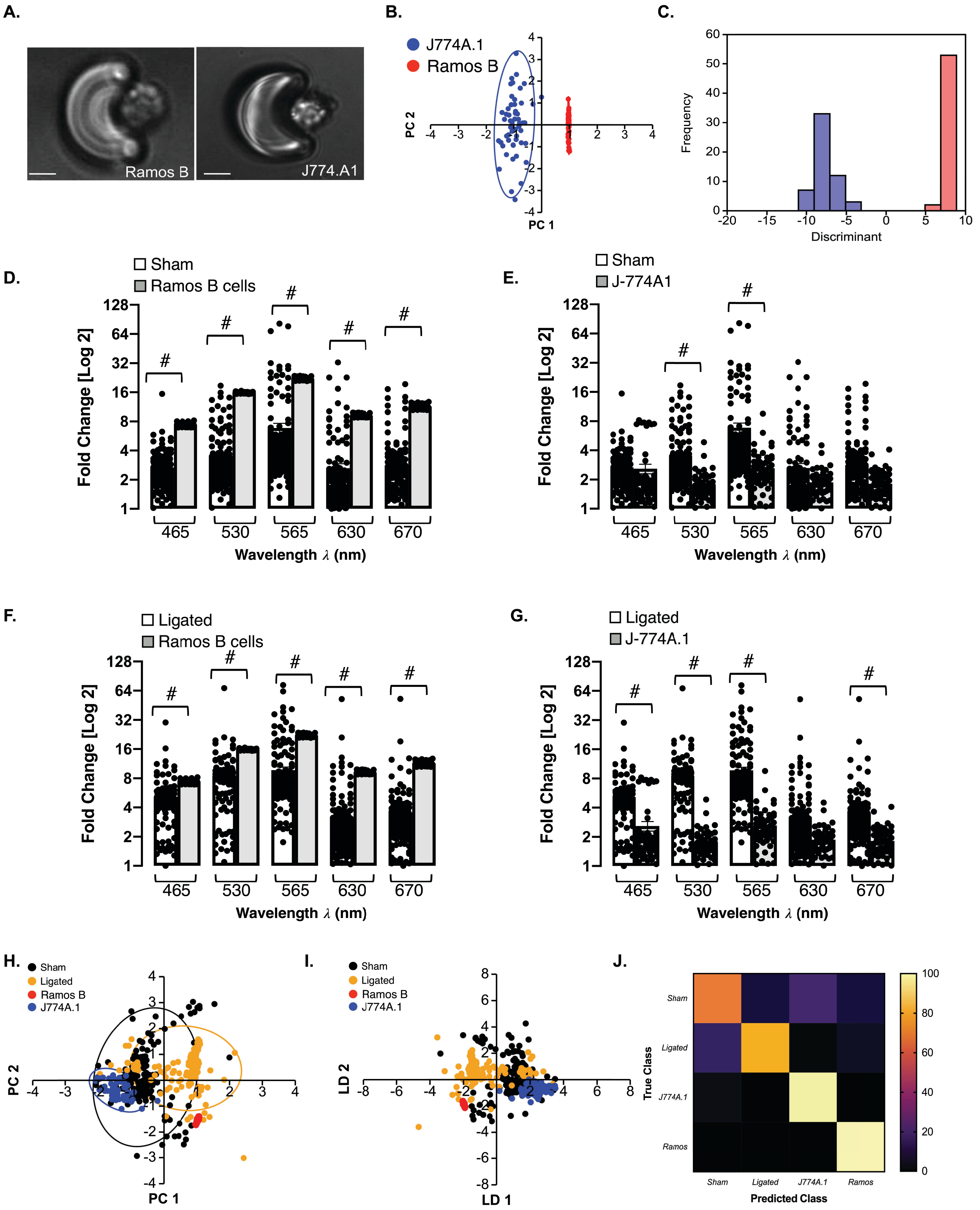

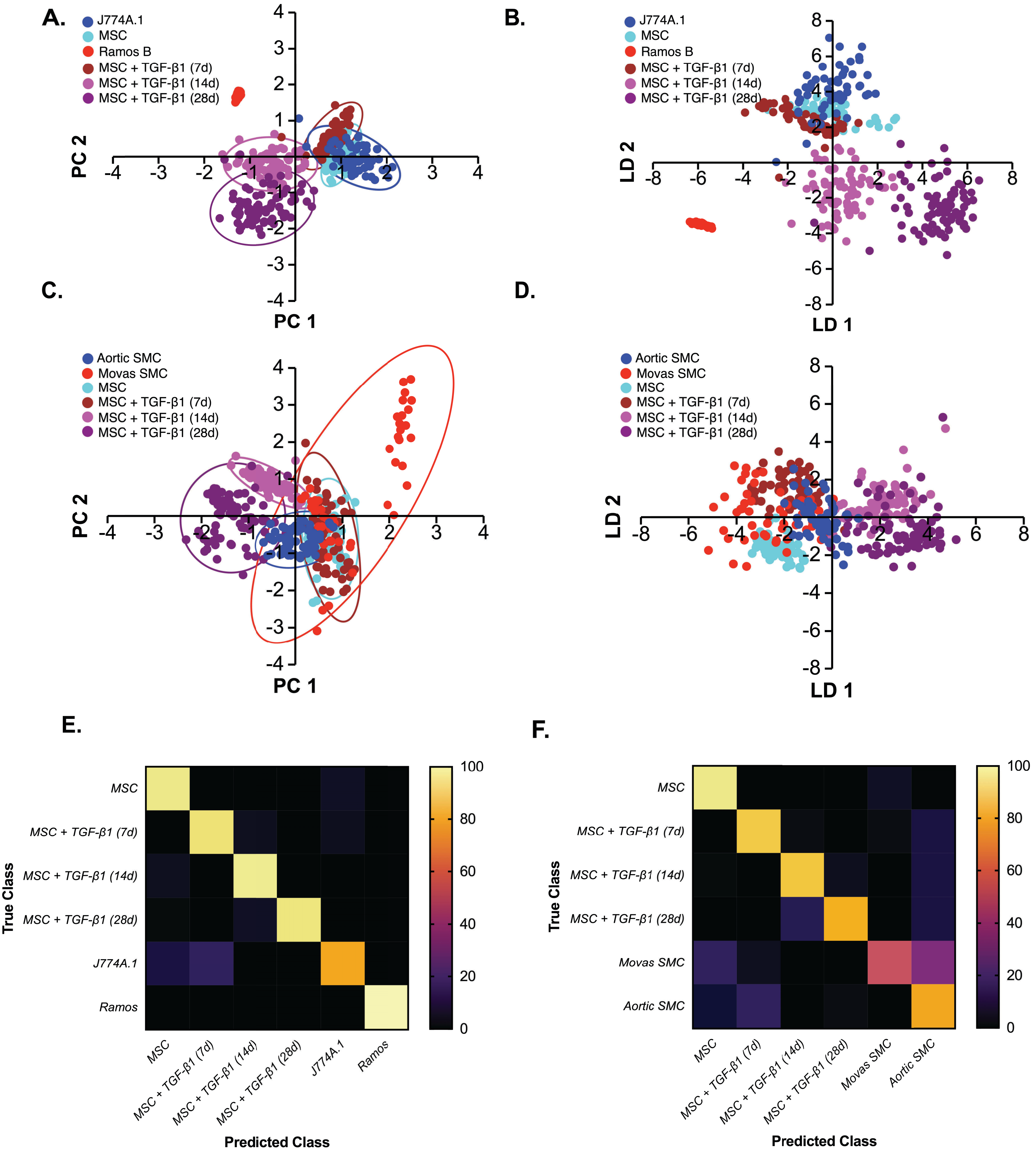

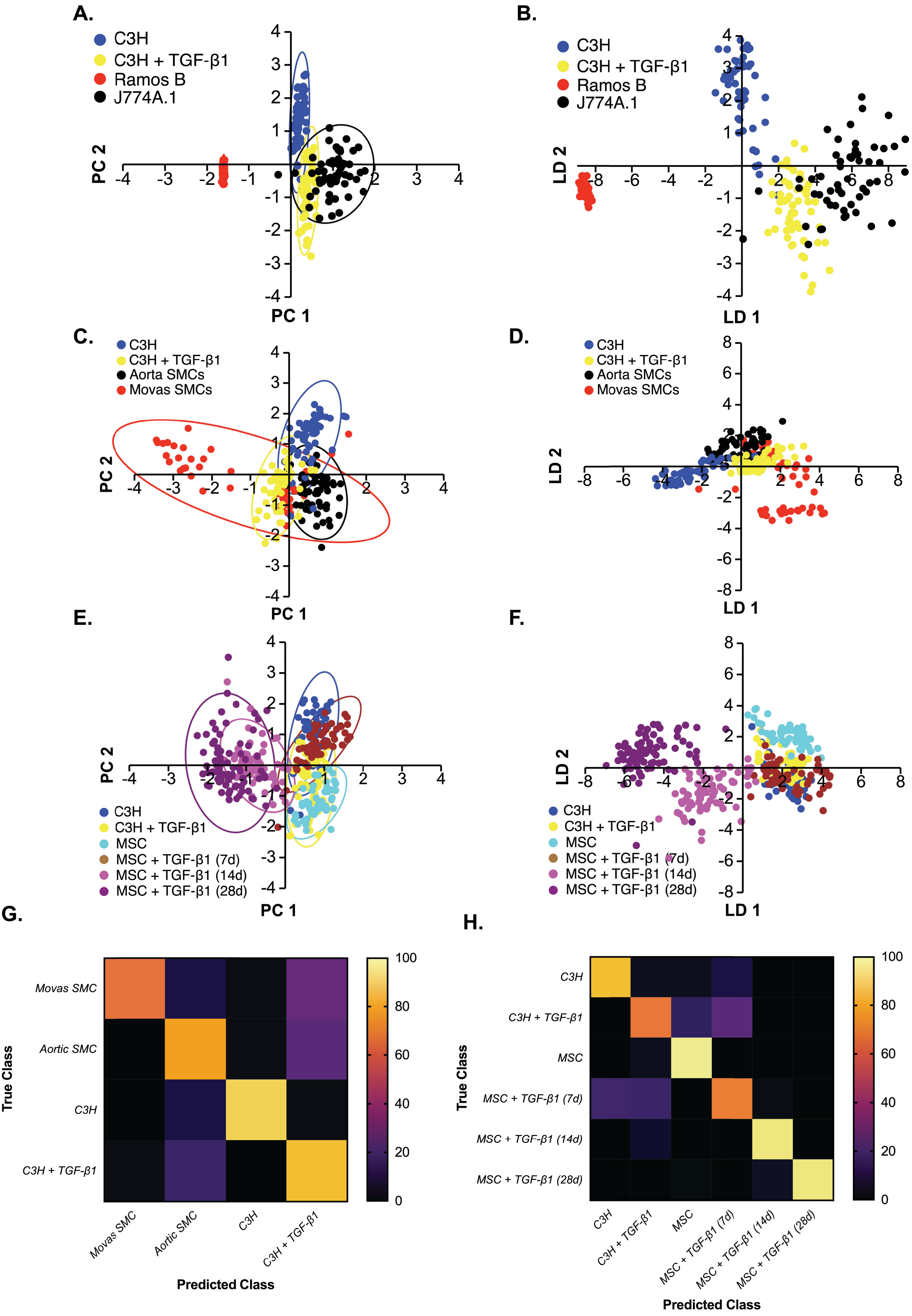

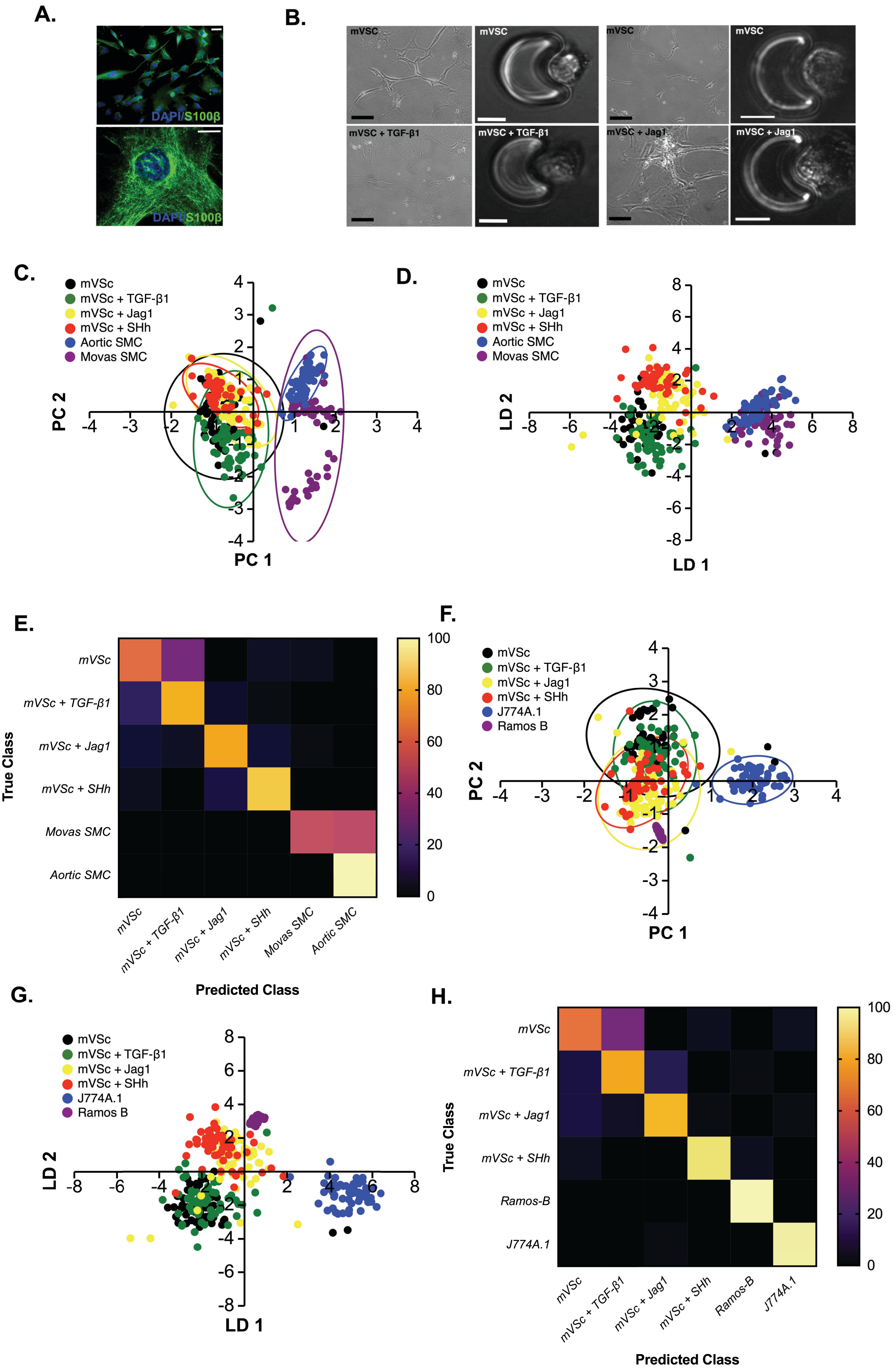

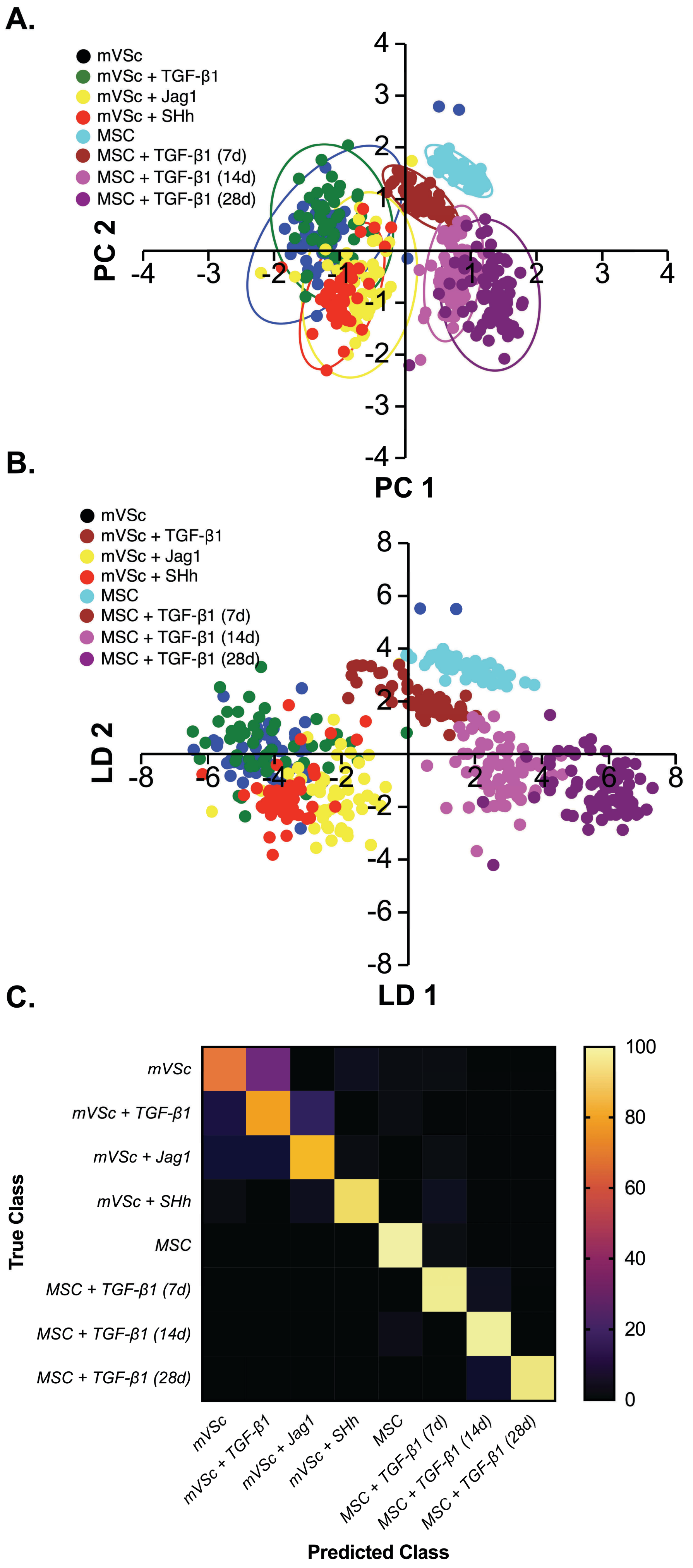

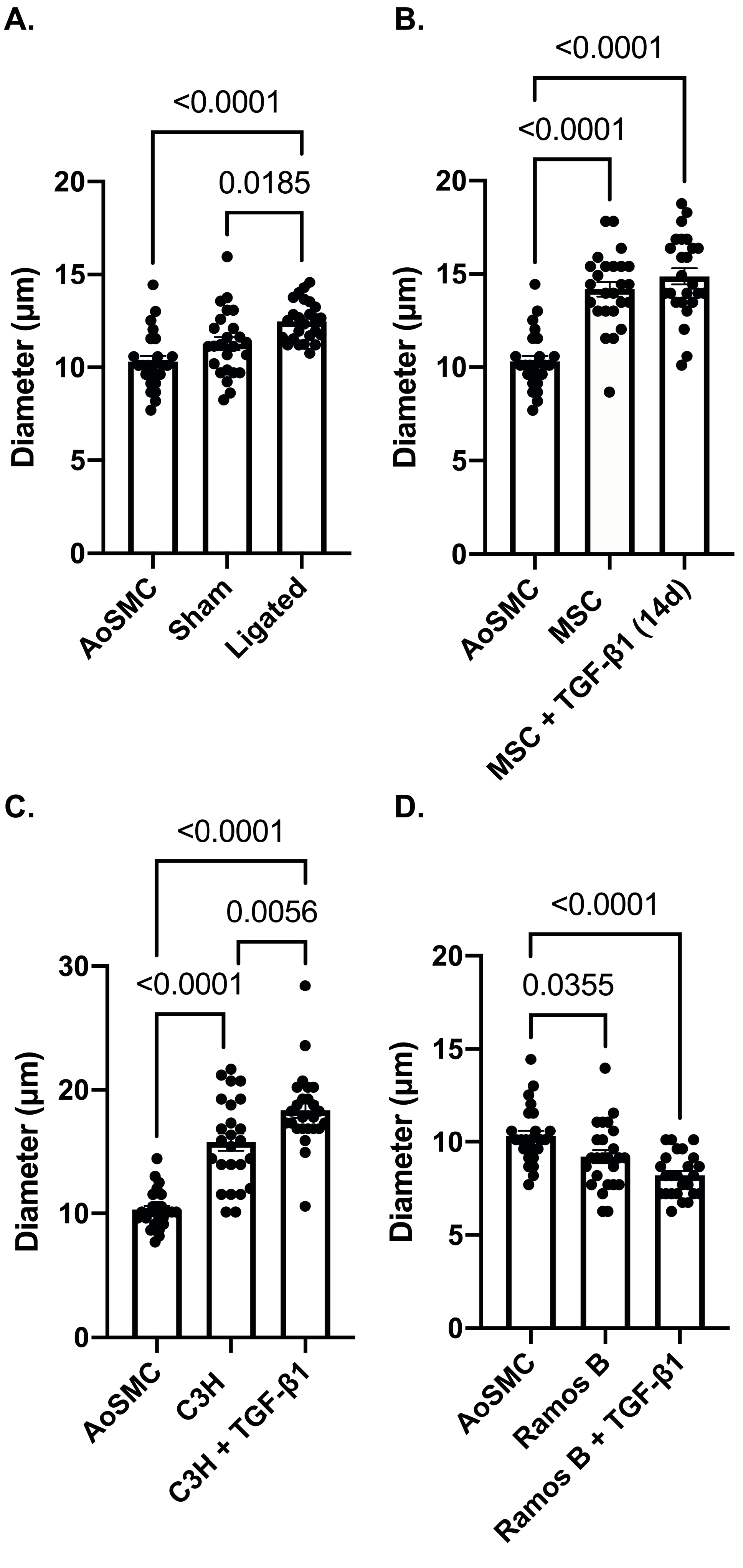

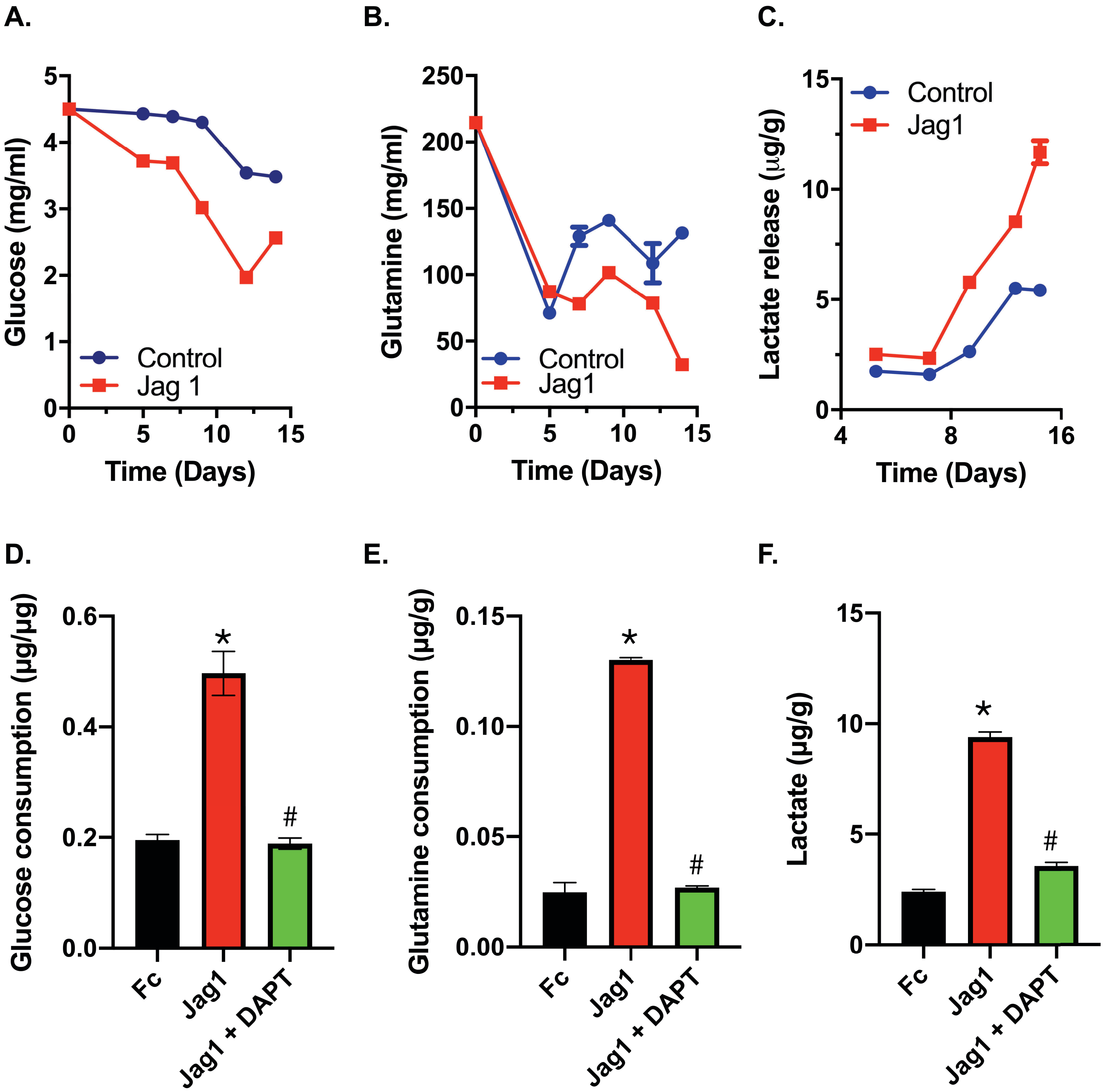

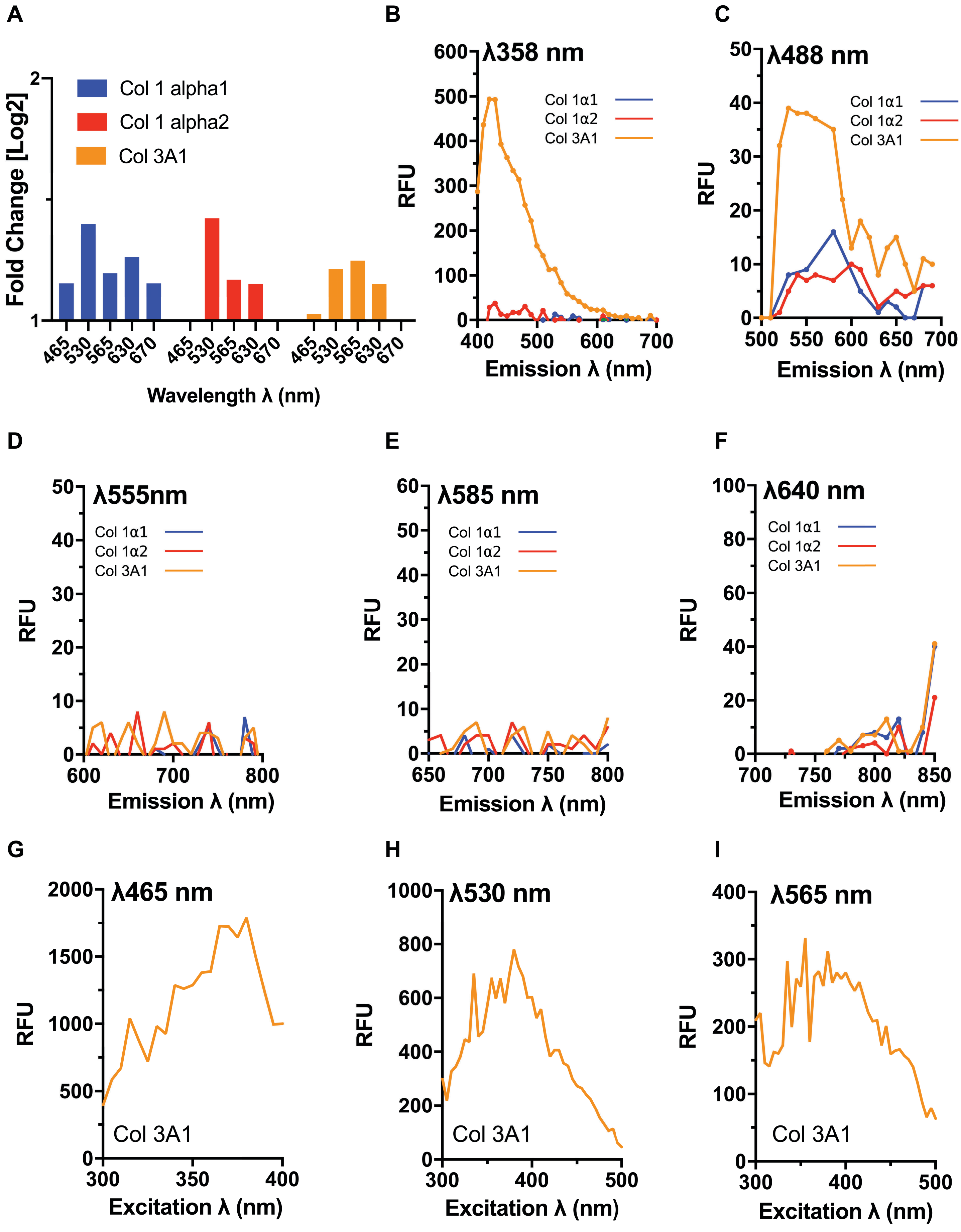
